# Transcriptomic expression patterns of two contrasting lowland rice varieties reveal high iron stress tolerance

**DOI:** 10.1101/2020.05.01.070516

**Authors:** Saradia Kar, Hans-Jörg Mai, Hadeel Khalouf, Heithem Ben Abdallah, Samantha Flachbart, Claudia Fink-Straube, Andrea Bräutigam, Guosheng Xiong, Lianguang Shang, Sanjib Kumar Panda, Petra Bauer

**Affiliations:** Institute of Botany, Heinrich Heine University, Universitätsstr. 1, D-40225 Düsseldorf, Germany; Plant Molecular Biotechnology Laboratory, Department of Life Science and Bioinformatics, Assam University, Silchar, India; Institute of Plant Biochemistry, Heinrich Heine University, Universitätsstr. 1, D-40225 Düsseldorf, Germany; Leibniz Institute for New Materials, D-66123 Saarbrücken, Germany; Faculty of Biology, Bielefeld University, Universitätsstr. 27, D-33615 Bielefeld; Agricultural Genomics Institute at Shenzhen, Chinese Academy of Agricultural Sciences, Shenzhen 518120, China; Cluster of Excellence on Plant Science (CEPLAS), Heinrich Heine University, Düsseldorf, Germany

**Author notes:** joint first co-authors (Saradia Kar, Hans-Jörg Mai). for correspondence: Petra Bauer. Author email addresses, Saradia Kar, Hans-Jörg Mai, Hadeel Khalouf, Heithem Ben Abdallah, Samantha Flachbart, Claudia Fink-Straube, Andrea Bräutigam, Guosheng Xiong, Lianguang Shang, Sanjib Kumar Panda, Petra Bauer.

**Keywords:** iron toxicity, iron uptake, leaf bronzing, metal homeostasis, *Oryza sativa*, oxidative stress, rice, RNA-Sequencing, susceptible, tolerant

## Abstract

Iron (Fe) toxicity is a major challenge for plant cultivation in acidic water-logged soil environments, where lowland rice is a major staple food crop. Only few studies addressed the molecular characterization of excess Fe tolerance in rice, and these highlight different mechanisms for Fe tolerance in the studied varieties.

Here, we screened 16 lowland rice varieties for excess Fe stress growth responses to identify contrasting lines, Fe-tolerant Lachit and -susceptible Hacha. Hacha and Lachit differed in their physiological and morphological responses to excess Fe, including leaf growth, leaf rolling, reactive oxygen species generation, Fe and metal contents. These responses were mirrored by differential gene expression patterns, obtained through RNA-sequencing, and corresponding GO term enrichment in tolerant versus susceptible lines. From the comparative transcriptomic profiles between Lachit and Hacha in response to excess Fe stress, individual genes of the category metal homeostasis, mainly root-expressed, may contribute to the tolerance of Lachit. 22 out of these 35 metal homeostasis genes are present in selection sweep genomic regions, in breeding signatures and/or differentiated during rice domestication. These findings will serve to design targeted Fe tolerance breeding of rice crops.

**Summary statement:** Lowland rice varieties Hacha and Lachit were selected for contrasting abilities to cope with iron excess stress. Morphological and physiological phenotypes were mirrored by molecular transcriptome changes, indicating altered metal homeostasis in the root as an adaptive tolerance mechanism in Lachit.

## Introduction

Iron (Fe) is a vital cofactor for many redox processes in plants. This micronutrient is needed during photosynthesis, respiration, nutrient assimilation and in many other metabolic pathways. Fe is very abundant in the soil. But since it is mostly bound to ferric hydroxides, Fe is normally not readily bio-available in the soil. Therefore, plants actively mobilize Fe, either by soil acidification, chelation by small organic molecules and reduction from ferric (Fe^3+^) to ferrous Fe (Fe^2+^) (Strategy I, found in most plant species) or by chelation of Fe^3+^ with phytosiderophores (Strategy II, in Poaceae). In acidic paddy field environments, Fe^3+^ is several orders of magnitude more soluble and more often reduced to Fe^2+^ than at neutral pH. Fe^2+^ is therefore bio-available, and plants can take it up without costly mobilization efforts.

Rice (*Oryza sativa*) is among the top most widely consumed staple foods (Das et al., 2010). More than 700 million tons of rice grains are produced on a yearly basis, about 90 % in Asia, less in Africa (Muthayya, Sugimoto, Montgomery, & Maberly, 2014). Despite of being a Strategy II phytosiderophore-producing crop, rice can acquire Fe directly as Fe^2+^ through the divalent metal Fe-regulated transporter of the root epidermis in paddy field conditions (Ishimaru et al., 2006). Fe toxicity is found under hypoxic conditions in Northeast India and in some parts of Africa. Northeast India has high potential for cultivation of diverse lowland rice varieties, however, its soils are strongly acidic with pH ranges from 5.5 to 5.1 (Barthakur, 2018). In this region, bio-available Fe is abundant leading to Fe toxicity as a major constraint affecting the annual yield (Baruah, Das, & Das, 2007).

A typical symptom of Fe toxicity is leaf bronzing with stress-induced anthocyanin pigmentation and necrotic lesions leading in severe cases even to plant failure (Tanaka, Loe, & Navasero, 1966). The toxicity of Fe is caused by oxidative stress. Fe^3+^ and Fe^2+^ stimulate generation of damaging hydroxyl and lipid alkoxyl radicals, through the Haber-Weiss and Fenton reactions. These radicals attack proteins, DNA and lipids, modify their reactive groups and cause decomposition of these biomolecules and mutations (Le, Brumbarova, & Bauer, 2019). Acidic soils are also associated with aluminum toxicity (Awasthi et al., 2017).

The molecular players and genetic background for Fe toxicity tolerance are not much understood. Few concrete breeding target loci are known and the mechanisms mostly not explained (J. Zhang et al., 2017). Analysis of genome-wide transcriptional response patterns revealed genotype-environment interactions and Fe tolerance response clusters and genes, which are partly at the origin of high Fe tolerance mechanisms described for rice at the morpho-physiological level. Several rice varieties tolerant to Fe toxicity exclude Fe, chelate or eliminate it, e.g. by repressing Fe acquisition in the root, forming an effective root plaque barrier to Fe and promoting root cell wall lignification, preventing Fe translocation to shoots, sequestering Fe in the cell wall and vacuole and/or chelating Fe by metal chelator nicotianamine (Aung et al., 2019; Stein et al., 2019; Tripathi et al., 2014; L. B. Wu et al., 2014). Exclusion and sequestration lower free cytoplasmic Fe levels, and are sometimes complemented by a second level of tolerance, which is to prevent cellular Fe and ROS toxicity through the action of antioxidants (Aung et al., 2019; Stein et al., 2019; Tripathi et al., 2014; L. B. Wu et al., 2014). Other effects may also occur, for example potassium transport may block root-to-shoot transport of Fe under high Fe stress (Diop et al., 2020; L.-B. Wu, Holtkamp, Wairich, & Frei, 2019). These studies show that different mechanisms may account for high Fe tolerance in different accessions of rice. Therefore, it is necessary to collect more data on the tolerance of individual rice varieties and correlate high Fe-associated gene expression to develop markers and deduce genetic players for high Fe tolerance.

Here, we screened 16 lowland rice varieties for high Fe tolerance in terms of root and shoot growth to identify contrasting lines, Fe-tolerant Lachit and susceptible Hacha. Lachit tolerated and survived Fe toxicity treatment in hydroponic culture in contrast to Hacha. Comparative transcriptomics via RNA-Sequencing (RNA-Seq) identified genes and physiological pathways associated with Fe toxicity tolerance in Lachit. These findings can be used to design and test targeted Fe tolerance breeding of rice crops.

## Materials and Methods

### Plant material

16 lowland *Oryza sativa* L. ssp. Indica varieties were obtained from the Regional Rice Research Station (RARS), Karimganj, Assam, India and the ICAR-NRRI-Regional Rainfed Lowland Rice Research Station (RRLRRS), Gerua, Guwahati, Assam, India. Hacha is high Fe-intolerant (derived from a cross of CRM 53 X IR 64), while Lachit is high Fe-tolerant (derived from a cross between CRM 13-3241 and Kalinga 2) (Awasthi et al., 2017).

### Plant growth

Plant growth was conducted in a controlled growth chamber, with a 16-hour light/ 8-hour dark cycle, 70 % humidity, a light intensity of 220 μmol*m^-2^*s^-1^, a day/night temperature of 26 °C. Rice seeds were disinfected using 1 % sodium hypochlorite for 15 min. Subsequently, seeds were soaked in water for one day at 4 °C and transferred for three days in the dark to 30 °C for germination. Homogenously grown seedlings were transplanted into 1 l pots filled with half-strength modified Hoagland solution for two days. Then, half-strength nutrient solution was replaced with a full-strength solution containing 1 mM calcium nitrate, 2.5 mM potassium nitrate, 2.86 mM ammonium nitrate, 1 mM potassium dihydrogen phosphate, 1 mM magnesium sulfate, 0.045 μM boric acid, 0.009 μM manganese chloride, 0.666 μM zinc sulfate, 0.4 μM copper sulfate, and 0.0176 μM sodium molybdate and 50 μM Fe sodium EDTA, pH 6.2, which also represented the control treatment condition. The nutrient solutions were renewed every second day. After another six days, a two-day Fe stress treatment with full-strength Hoagland medium plus 15 mM Fe sulfate at pH 5.2 was applied or plants were exposed to control conditions.

For revival experiments, the plants were re-exposed to control medium after the two-day stress condition and again the medium was renewed every second day. For the RNA-Seq experiment, eight plants were pooled for a biological sample. In total, three biological replicates were prepared. The revival experiment was conducted with eight plants per condition. For physiological and morphological experiments three to eight plants were grown as indicated in the figure legends.

### Morphological examination of plants

The shoot dry weight (SDW) was determined after dissecting the root system and fully drying the shoot at 120°C. The relative decrease in SDW (RDSDW) score was calculated in % as SDW_control-excess_*100/SDW_control_. Leaf lengths were measured from the base of the sheath to the tip of the blade. Leaves were dissected, weighed to determine the fresh weight, and after fully drying at 120°C to determine the leaf dry weight. Leaf rolling was assessed as weak, medium or strong as indicated in Fig. 3A.

### Hydrogen peroxide (H_2_O_2_) detection

Diaminobenzidine (DAB) staining was used to visualize H_2_O_2_ *in vivo*. L2 leaf blades were excised, vacuum-infiltrated for five minutes in a solution of 1 mg/ml 3,3’-DAB tetrahydrochloride in 50 mM MES, pH 6.5, followed by a one-hour incubation in a shaker and washed with distilled water. The sample was placed into 80 % ethanol and boiled at 80 °C until the leaves were de-stained from chlorophyll. The reaction was stopped by washing three times with distilled water. The samples were observed using a microscope.

H_2_O_2_ contents were quantified using the Amplex Red kit (Thermo Fisher Scientific). 50 mg plant sample were ground in 200 μl potassium phosphate buffer and 50 μl supernatant were used for horseradish peroxidase-based Amplex Red assay reactions, using appropriate controls, as described (T. Brumbarova, Le, & Bauer, 2016). The absorbance at 560 nm was used to calculate the content of H_2_O_2_ based on a mass standard curve.

### Malondialdehyde content

Lipid peroxidation was determined using colorimetric detection via the thiobarbituric acid (TBA)-malondialdehyde (MDA) assay, according to (Z. Zhang & Huang, 2013). Briefly, 50 mg of plant material was homogenized with 2 ml trichloroacetic acid (TCA, 0.1 %). After centrifugation, 1 ml supernatant was mixed with 2 ml of 0.5 % (w/v) TBA in 20 % (w/v) TCA and 10 μl of 4 % butylated hydroxytoluene. After 30-minute boiling and cooling the absorbance was determined at 532 and 600 nm. MDA concentrations were calculated after subtracting OD_532_ – OD_600_ and using the molar extinction coefficient of MDA (155 mM^-1^ * cm^-1^).

### Determination of metal contents and Perls Fe staining

L2 leaves and root systems were air-dried and dried at 100 °C for ten hours, finely powdered with an agate mortar and pestle. Metal contents (Fe, Zn, Mn, Cu) were determined following microwave digestion of dried samples with nitric acid and H_2_O_2_ (Multiwave 3000, Anton Paar) by high-resolution continuum source atomic absorption spectrometry HR-CS AAS (contrAA 700, Analytik Jena AG) at the Leibniz Institute for New Materials (INM, Saarbrücken) and calculated using reference metal contents

Perls Fe staining was conducted with 50-100 μm hand-made sections of L2 leaf sheaths and roots, according to (Tzvetina Brumbarova & Ivanov, 2014). Briefly, sections were de-stained, and then subjected to Perls Prussian Blue staining with potassium ferrocyanide solution to yield a blue precipitate. 50-100 μm hand-made cross-sections were prepared. For general anatomy observations, the sections were incubated for 5-8 minutes in 1 mg/ml fuchsine, 1 mg/ml chrysoidine, 1 mg/ml astra blue, 0.025 ml/ml acetic acid staining (= FCA staining) solution. Sections were washed in distilled water, twice in 30 % ethanol, 70 % ethanol, again twice 30 % ethanol and isopropanol and mounted in distilled water on slides.

### RNA isolation and RNA sequencing (RNA-seq)

L2 leaves and the entire root systems from eight plants per sample were dissected from Hacha and Lachit, exposed to a control growth condition or two-day 15 mM Fe excess condition. Three independent biological replicates were harvested per experimental condition and line. RNA was isolated from 100 mg plant material using the RNeasy plant mini kit (Qiagen). RNA was checked and sequenced according to (Brilhaus, Bräutigam, Mettler-Altmann, Winter, & Weber, 2016). Briefly, the quality and quantity of RNA was analyzed following DNase digestion using the Bioanalyzer 2100 (Agilent Technologies). RNA samples with intact bands and an RNA integrity number (RIN) >7.0 were used for preparation of RNA-seq libraries with TruSeq RNA Sample Prep Kit v2 (Illumina Technologies). Double-stranded cDNA was purified for end repair, adaptor ligation, and DNA fragment enrichment. The libraries were sequenced as 151 bp paired-end reads using Illumina HiSeq at the Biologisch-Medizinisches Forschungszentrum, Heinrich Heine University Dusseldorf. Basecalling and demultiplexing was performed using bcl2fastq2. RNAseq data are deposited in GEO.

### Analysis of RNA-Seq data and of candidate genes

Paired reads were trimmed with trimmomatic (Bolger, Lohse, & Usadel, 2014) and their quality evaluated with fastqc (Andrews, 2010). Using kallisto (Bray, Pimentel, Melsted, & Pachter, 2016), the trimmed reads were mapped to the *Oryza sativa* transcriptome assembly (Oryza_sativa.IRGSP-1.0.cdna.all.fa, downloaded from EnsemblPlants) and quantified. *Arabidopsis thaliana* homologs were determined by blasting the rice transcriptome against the *A. thaliana* proteome using the rice CDS collection (Oryza_sativa.IRGSP-1.0.cds.all.fa; downloaded from Gramene) and *A. thaliana* proteome sequences (TAIR10_pep_20101214_updated; downloaded from TAIR), using the blastx algorithm of the Blast+ suite (Altschul, Gish, Miller, Myers, & Lipman, 1990) and genes were called similar if the e-value was < 1E-5. Among multiple hits, Arabidopsis proteins with highest score followed by lowest E value were taken. Transcripts per million (TPM) counts for all samples were used for principal component analyses (PCA, R: prcomp), hierarchical clustering (HC, R: dist, hclust) and plots to visualize biological replicate quality. Estimated counts of all replicates were statistically analyzed using edgeR (McCarthy, Chen, & Smyth, 2012; Robinson, McCarthy, & Smyth, 2010). Resulting p-values were adjusted with the Bonferroni method. Fold-change values of gene expression were calculated in pairwise comparisons between Lachit and Hacha upon Fe excess stress or the control for roots and leaves, and within a line between excess Fe stress and the control for roots and leaves (in total eight meaningful comparisons). All assembled data including absolute expression data, fold-change ratios, p-values and annotations are provided in Supplemental Table S1. Genes were considered differentially expressed if their adjusted p-value was < 0.01 in at least one pair-wise comparison (Supplemental Table S2). Gene ontology (GO) analysis was carried out using topGO (Alexa & Rahnenfuhrer, 2010) with the rice GO annotations (go_ensembl_oryza_sativa.gaf; downloaded from Gramene), applying Fisher’s exact test.

For analysis of selection sweep regions, gene loci were compared to selection sweep regions and breeding signatures as described (Huang et al., 2012; Xie et al., 2015). We also performed diversity statistics (π), the population-differentiation statistics (Fst), and crosspopulation likelihood (XP-CLR) analysis using 147 rice accession as described (Wang et al., 2020). Candidate targets were determined from SNP haplotype analysis and phylogenetic trees were constructed by FastTree software (Price, Dehal, & Arkin, 2010) using selected 230 rice accessions in a representative mini core collection (MCC) constructed by (H. Zhang et al., 2011).

### Gene expression by reverse transcription-quantitative PCR (RT-qPCR)

RT-qPCR was conducted with RNA samples prepared for RNA-seq using three biological replicates for each sample. cDNA synthesis and qPCR were carried out as described previously (Ben Abdallah & Bauer, 2016). Briefly, total RNA was used for cDNA synthesis using an oligo-dT primer. qPCR was conducted using the SYBR Green detection method. Each biological cDNA sample was tested in two technical qPCR replicates. Absolute starting quantities of templates were determined by mass standard curve analysis. Normalized absolute expression data were obtained after normalization to the internal reference genes. The primer sequences (5’ to 3’ direction) for reference genes were *OsTUB1* CTCATCAGTGGCAAGGAGGA and CTCCAACAGCGTTGAACACA (OS11G0247300, tubulin 1 gene), *OsUBI1* CTTCACCTTGTCCTCCGTCT and TTGTCCTGGATCTTGGCCTT (Os02G0161900, ubiquitin 1 gene). The primer sequences (5’ to 3’ direction) for genes of interest were *OsFER1* TCTTCCAGCCATTCGAGGAG and TGACTCCATGCATAGCCACA (OS11G0106700, ferritin 1 gene), *OsMT1g, OsMTP8* ACTTGCCCTCTACATATACTGCA and TCACTCAGACATACAGATTCCTC (OS02G0775100, metal tolerance protein 8 gene), *OsPRX131* GCTAGATGACACCCCAACCT and AGGGAAGTGTCGATGTTGGT (Os11g0112200, peroxidase 131 gene), *OsSALT* AGGCGTGACAATCTACAGCT and ATCATAGACTGGGCCATGGG (Os01g0348900, *SALT STRESS-INDUCED* gene), *OsASR3* CAAGAAGGACGGGGAGGAG AND CACCTCCTCCTTCACCTTGT (OS02G33820, *ABIOTIC STRESS-RESPONSIVE* GENE), *OsNRAMP4* ACTGTGCAGGGAAGAGTACC and GGGGTCCAACTTTCTTGCTG (OS02G0131800, *OsNRAMP4* metal transporter gene), *OsYSL15* GCAACACAGCATCAGAGGAG and CCAGGCGTAAACTTCATGCA (OS02G0650300, *OsYSL15* phytosiderophore importer gene).

### Statistical Analysis

Statistical analyses of morphological, physiological and RT-qPCR gene expression data were carried out using R. Comparison of means were performed using one-way analysis of variance (ANOVA) and Tukey honest significant difference (HSD) test. RNA-Seq data were analyzed as described above and using exact tests for differences between two groups of negative-binomial counts (Robinson & Smyth, 2007) with Bonferroni correction.

## Results

### Lowland Indica rice varieties Hacha and Lachit have contrasting abilities to deal with excess Fe stress

16 lowland *Oryza sativa* Indica varieties from Northeast India were screened under high Fe and control conditions for their relative decrease in shoot dry weight (RDSDW) upon excess Fe stress. Lachit had the lowest RDSDW value, indicative of best tolerance, while Hacha had the highest value and was most susceptible among the tested lines (Figure 1). Lachit and Hacha were chosen as contrasting pair. Plants of the two lines were exposed to 48-hour excess Fe stress and control treatments and their response patterns recorded. The Fe stress consisted of 15 mM Fe sulfate treatment at pH 5.2 versus regular 50 μM Fe at pH 6.2 control condition. Recovery from the excess Fe stress was tested in a revival experiment that lasted up to 6 days following the treatments, when plants were re-exposed back to regular control conditions (Figure 2). Under control treatment, Hacha and Lachit L2 and L3 leaves increased in size steadily for 1 −1.3 cm during the 6 days (Figure 2). L3 leaves grew more than L2 leaves. Hacha L3 leaves gained an average of 2.5 ±0.49 cm (value ±SD), and Lachit L3 leaves gained nearly 6 ±1.6 cm during this time in the control condition, compared with 0.9 ±0.8 cm and 1.3 ±0.6 cm in the L2 leaves, respectively (Figure 2B, 2D compared to Figure 2A, 2C). Following excess Fe stress, Hacha L2 and L3 leaves did not grow further during the 6 days (Figure 2). Lachit, on the other hand, was able to recover and to survive the Fe stress. Lachit L2 and L3 leaves grew 0.5 and 0.4 cm, respectively, during the 6 day recovery period after the stress, but did not reach the sizes of control leaves (Figure 2). Even though the length increases in Lachit leaves were not statistically different from those of Hacha leaves (Figure 2), Lachit plants had grown further and ultimately had survived, in contrast to Hacha. Thus, based on their growth phenotypes, Lachit plants tolerated the high Fe treatment but not Hacha plants.

**Figure 1.**
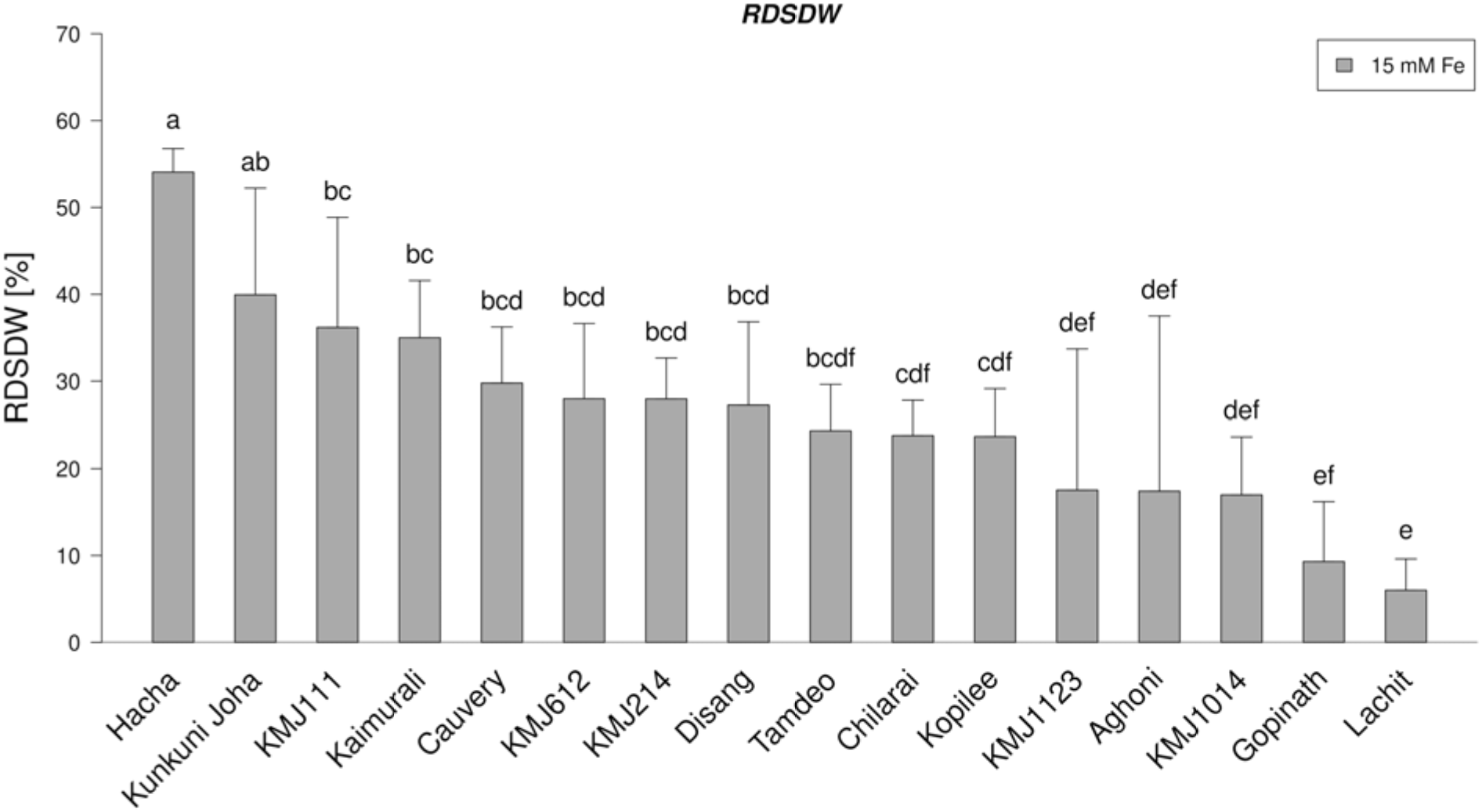
Selection of two rice varieties with contrasting abilities to sustain growth under Fe excess stress using the relative decrease in shoot dry weight (RDSDW). Different letters, indicative of significant difference (p < 0.05; n = 9; ANOVA with Tukey HSD).

**Figure 2.**
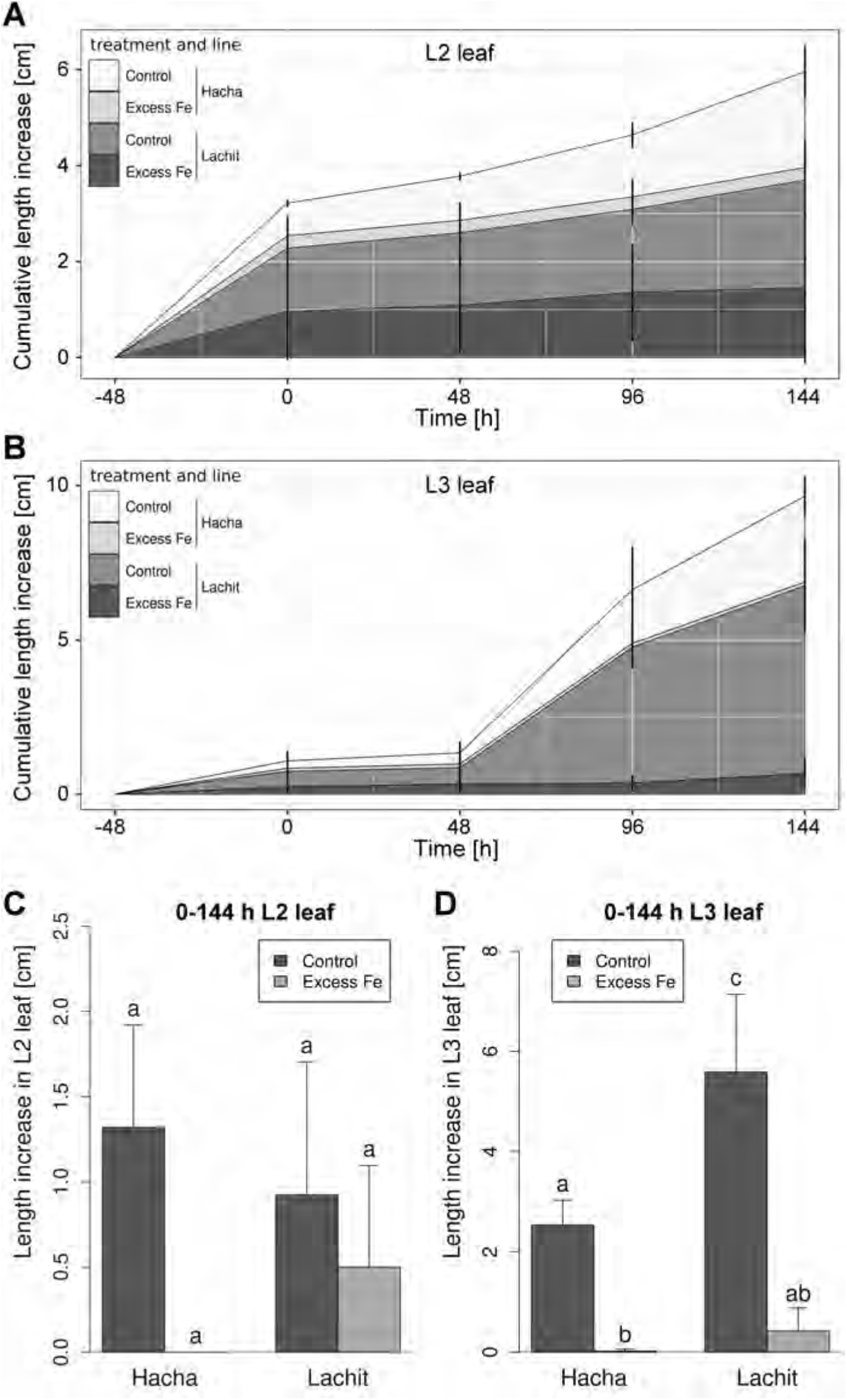
Excess Fe stress and revival experiment. (A, B) Stacked area plots of the cumulative length increases of (A), L2 and (B), L3 leaves in Hacha and Lachit, recorded from the beginning of excess Fe stress (−48 h), the end of Fe excess stress (0 h) and during a revival phase after the end of the stress period (48, 96 and 144 h) and respective controls. (C, D) Length increases of (C), L2 leaves; (D), L3 leaves, between 0 h and 144 h after the end of the excess iron and control treatments. Different letters indicate a significant difference between the respective groups (p < 0.05; n = 3; ANOVA with Tukey HSD).

Fe excess caused leaf rolling (Figure 3A). After two days of excess Fe stress, Hacha showed stronger leaf rolling symptoms than Lachit (Figure 3B). Leaf rolling might be coupled to drought. Therefore, we assessed fresh to dry weight ratios to possibly link dehydration and high Fe stress. The fresh to dry weight ratios were lower upon excess Fe treatment than the control in both lines, however, the ratios remained at comparable numbers in Lachit and Hacha (Figure 3C). Taken together, leaf rolling was a symptom of high Fe stress, more pronounced in Hacha than Lachit. Despite of that, different degrees of leaf rolling had minor effects on the overall fresh to dry weight ratios.

High Fe causes oxidative stress in cells with formation of reactive oxygen species (ROS), such as hydrogen peroxide (H_2_O_2_) and lipid peroxide. H_2_O_2_ generation was qualitatively determined by diaminobenzidine (DAB) staining. Excess Fe-stressed plants showed stronger DAB staining in leaves of both lines than controls (Figure 4A). Quantitative measurements of H_2_O_2_ confirmed higher levels in leaves upon Fe excess versus the controls, but did not uncover differences between the lines (Figure 4B). Malondialdehyde (MDA) is a byproduct of polyunsaturated fatty acid breakdown, occurring upon lipid peroxidation, catalyzed by Fe. Contents of MDA were elevated upon Fe excess stress in leaves of both varieties, whereby MDA contents were lower under control and Fe excess stress conditions in Lachit compared with Hacha (Figure 4C). Taken together, ROS generation is coupled to high Fe, and this effect is stronger for MDA generation in Hacha than Lachit.

**Figure 3.**
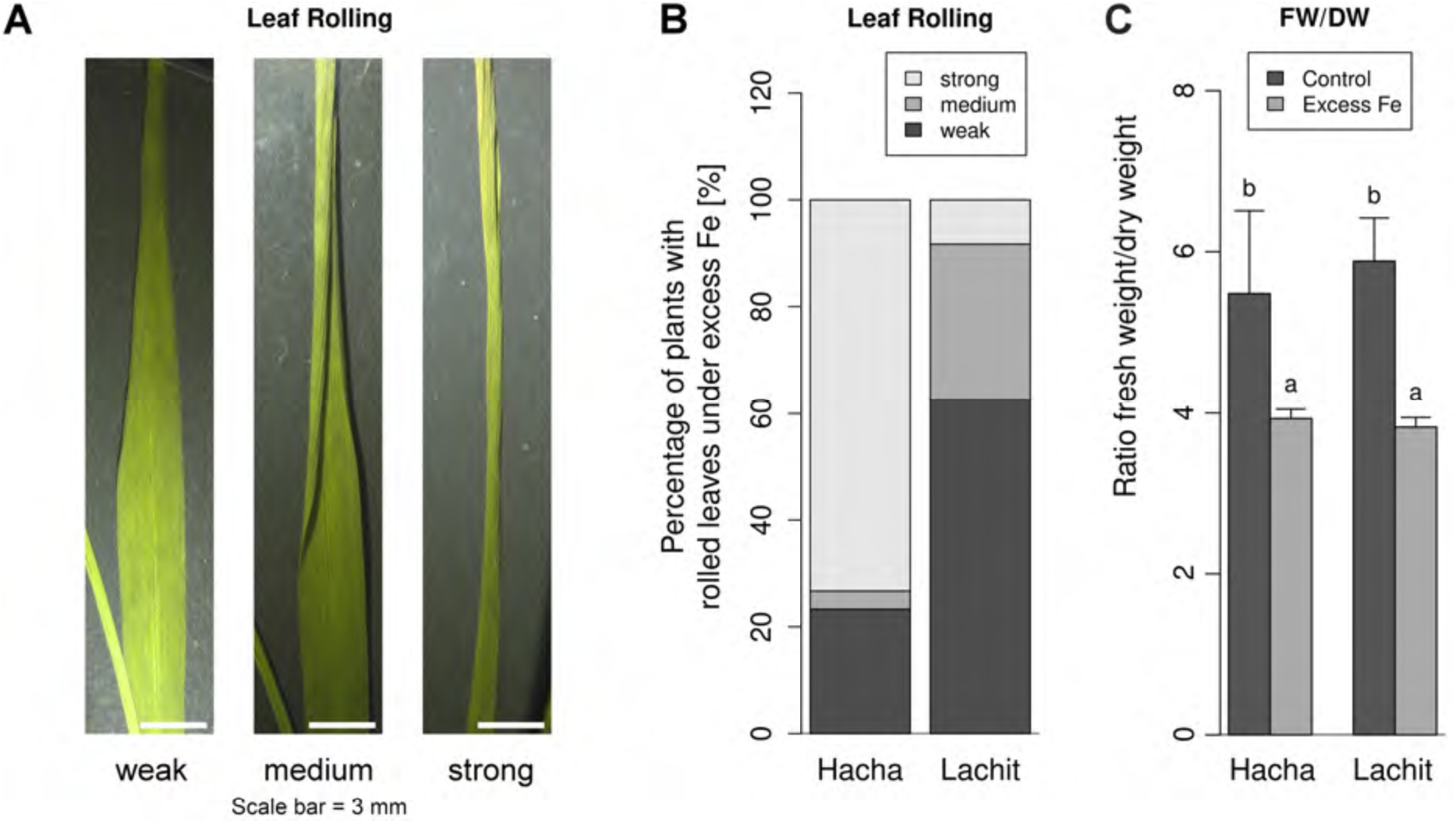
Leaf rolling and fresh to dry weight ratios. (A, B) Leaf rolling symptoms of Fe excess stress with (A), photos and scale of weak to strong leaf rolling phenotypes; (B), proportions of weak, medium and strong leaf rolling symptoms in Hacha and Lachit exposed to excess Fe stress. (C) Fresh weight (FW) to dry weight (DW) ratios. Different letters indicate significant differences between samples (p < 0.05; n = 3; ANOVA with Tukey HSD).

**Figure 4.**
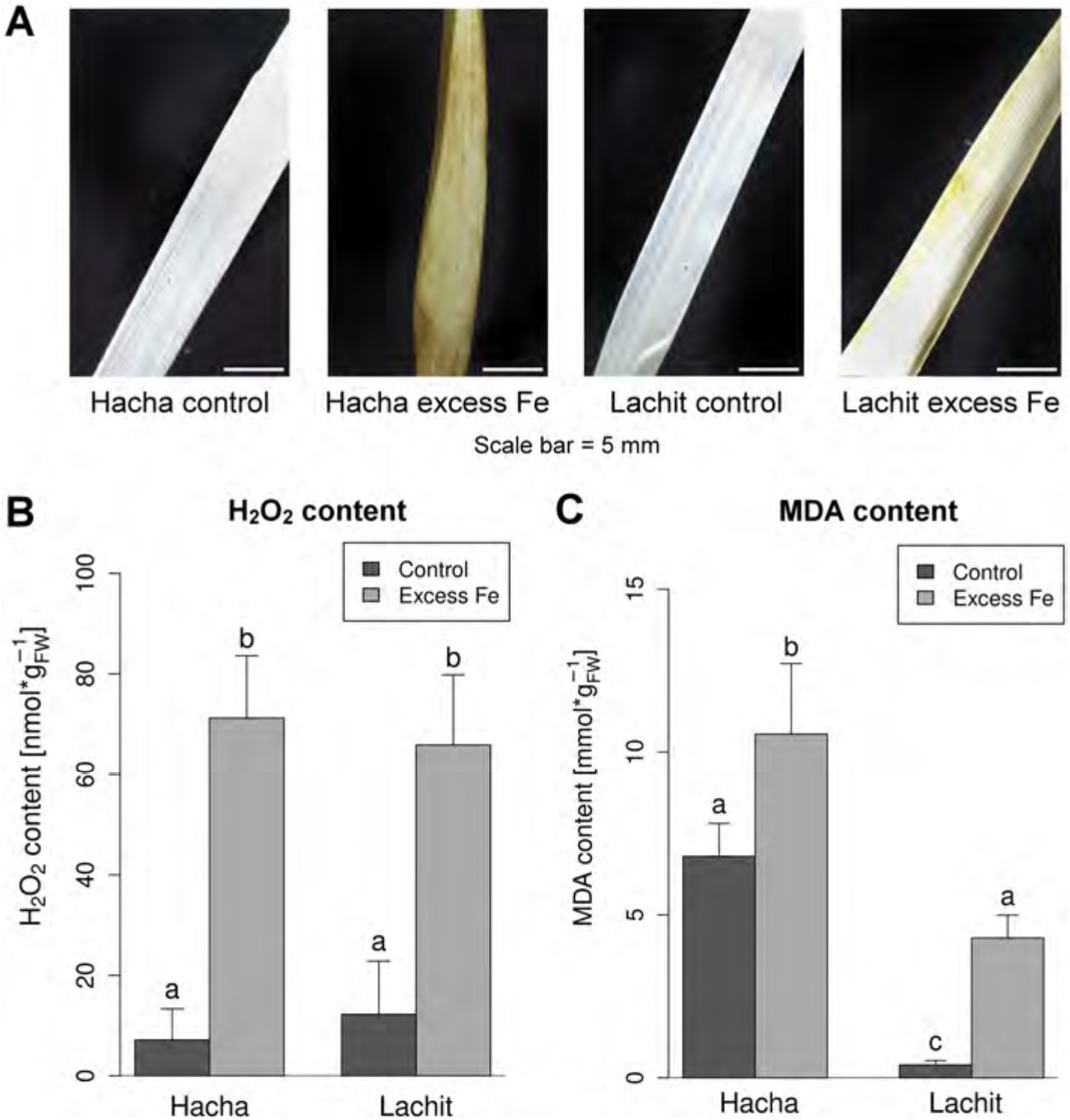
Oxidative stress under Fe excess in L2 leaves. (A), Photos of diaminobenzidine (DAB)-stained leaves; (B), hydrogen peroxide (H_2_O_2_) contents; (C), malondialdehyde (MDA) contents. Different letters, significant differences between samples (p < 0.05; n = 3; ANOVA with Tukey HSD).

The reactions of leaves to high Fe treatment indicate that excessive Fe must have been transported from roots to leaves to some degree in both genotypes. Under control conditions, Hacha and Lachit had similar Fe, zinc (Zn), manganese (Mn) and copper (Cu) contents (Figure 5A–5D). As expected, Fe-stressed Hacha leaves had an almost 300 times higher Fe content than control leaves and Fe-stressed Lachit leaves had about 200 times higher Fe contents than control leaves (Figure 5A). Lachit leaves had lower Fe levels than Hacha plants, in response to Fe excess (Figure 5A). Under excess Fe, Lachit had also lower levels of Zn, Mn and Cu contents compared with Hacha (Figure 5B-5D). Cu contents were slightly increased in Hacha under excess Fe compared to the control (Figure 5D). Taken together, metal content measurements strongly suggest that lower contents of all four metals in leaves of Lachit contributed to the lower metal toxicity phenotype observed in this line.

**Figure 5.**
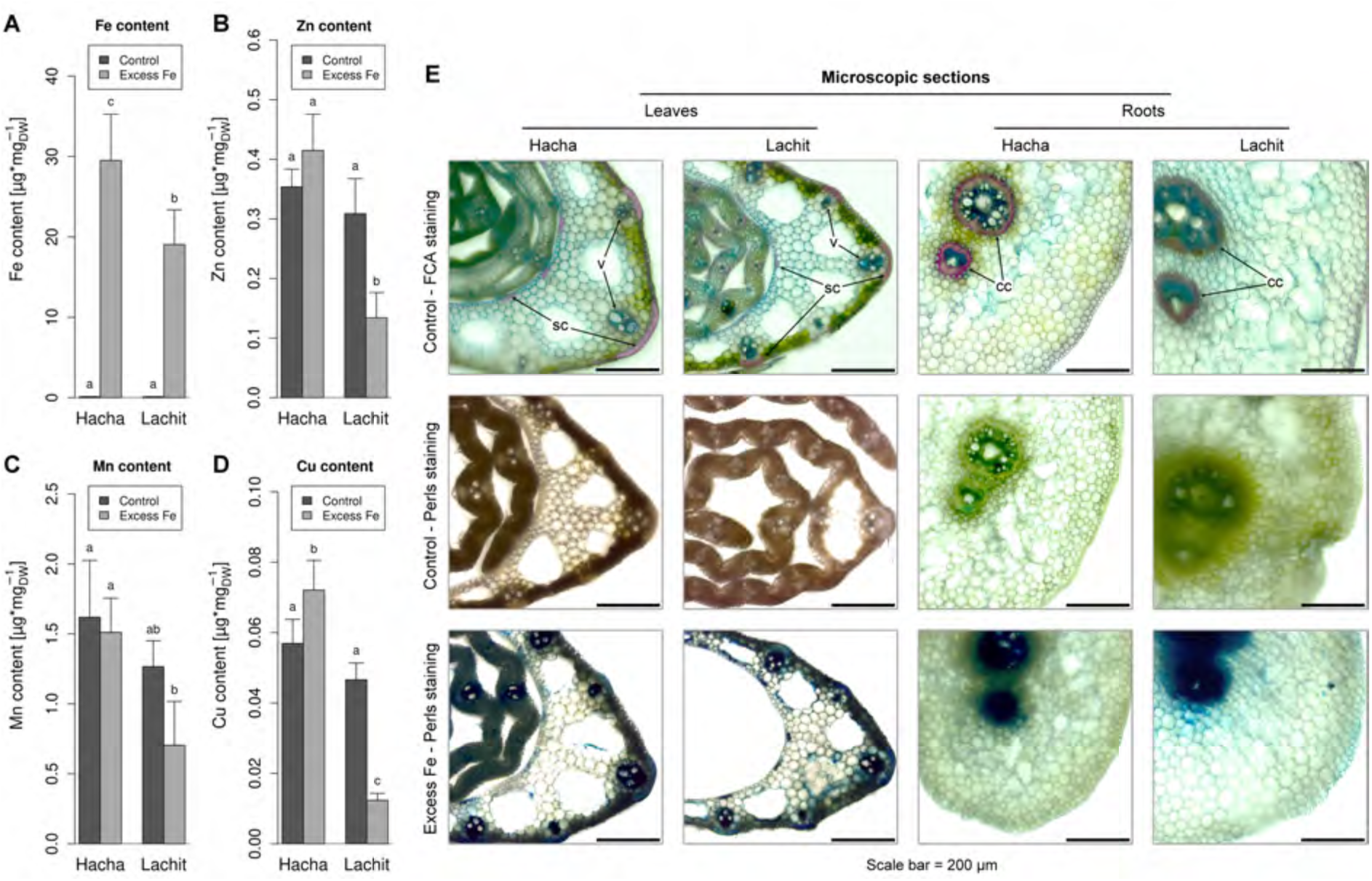
Metal contents and Fe distribution. (A-D), Contents of (A), Fe; (B), Zn; (C), Mn; (D), Cu contents per dry weight (DW) in L2 leaves of Hacha and Lachit in response to excess Fe and control. Different letters, significant differences between samples (p < 0.05; n = 3; ANOVA with Tukey HSD). (E), Microscopic images of cross-sections of Hacha and Lachit L2 leaves and roots following FCA-staining (general anatomy) and Perls-staining (Fe deposits); v, veins (leaves); sc (leaves), sclerenchyma cell rows; and cc, central cylinder (roots) are designated by arrowheads. Scale bar 200 μm.

Fe deposits can be localized in distinct tissues via Perls Fe staining. Hacha and Lachit did not differ in their Fe localization patterns in leaves and roots (Figure 5E). Fe deposits were found in parenchyma cells, that have photosynthetic functions, in vascular bundles, and in sclerenchyma cell rows present on either side of the veins underneath the epidermis. In roots, Fe was deposited in cells of the central cylinder (Figure 5E).

In summary, Lachit tolerated the Fe stress condition, leaves grew more and plants survived the stress, in contrast to Hacha. Hacha was more susceptible to Fe excess stress and had enhanced Fe levels compared with Lachit. Fe tolerance in Lachit can be explained by a lower uptake or translocation of Fe, Zn, Mn and Cu from root to shoot in Lachit, and consequently resulting in less oxidative stress seen as lipid peroxidation.

### Comparative RNA-Seq transcriptome analysis of Lachit and Hacha uncovers key regulatory patterns to excess Fe stress

Comparative transcriptomics via RNA-Seq is an approach to uncover biological mechanisms based on gene expression pattern differences between genetic variants and treatments. Hacha and Lachit were exposed to the two-day excess Fe stress and control conditions. Roots and L2 leaves were harvested, pooled from eight plants per biological replicate and used for the RNA-Seq analysis pipeline, with a total of three biological replicates (Supplemental Figure S1). Based on the excess Fe phenotype in the leaves of both genotypes we expected strong transcriptional changes in leaves with the response being larger in Hacha. We also expected differences between Lachit and Hacha in roots some of which may explain Lachit’s resistance to excess Fe, e.g. explaining lower uptake or translocation of Fe to shoots.

Principle component analysis (PCA) and hierarchical clustering (HC) served to control the quality of samples and assess the different responses of Hacha and Lachit. PCA separates data according to their variation with descending significance of each subsequent principal component. The most significant principal component (PC1) separated the tissue types, roots versus leaves, with 52 % of the variation (Figure 6A). The second principal component (PC2) separated Fe excess treatment versus control with 22 % of the variation (Figure 6A). The third principal component (PC3) resolved cultivar-specific differences, which accounted for 7 % of the variation (Figure 6B). For both, leaf and root samples, the Fe effect was different between lines. The excess Fe-treated Lachit samples were closer to their controls than the excess Fe-treated Hacha samples (Figure 6A). Hierarchical clustering confirmed PCA results. HC resolved the same three parameters in the same order of significance (see positions of bifurcations in Supplemental Figure S2A).

**Figure 6.**
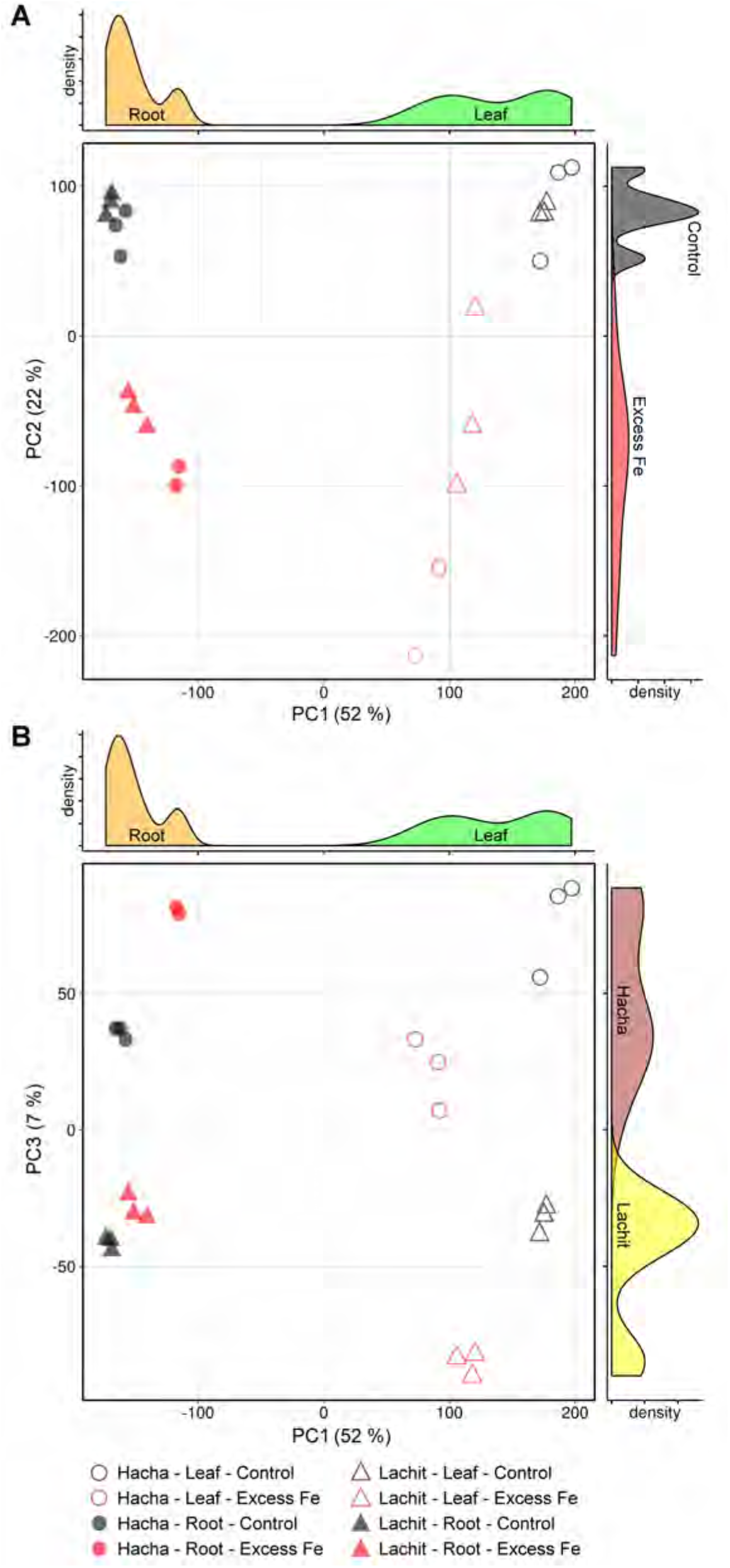
Principal component analysis (PCA) biplots of RNA-Seq samples and marginal density plots. The marginal density plots serve to illustrate the separation between the respective plant lines, organs or treatments in the biplots. (A), Principal component (PC) 1 vs. PC2; (B), PC1 vs. PC3. PC1, separation of roots and leaves; PC2, separation of excess Fe and control; PC3, separation of Hacha and Lachit.

To test whether one or both genotypes show excess Fe responses, differential transcript accumulation in roots and shoots in both genotypes was assessed (Figure 7A). Overall, more changes were observed in roots compared to shoots (Figure 7A). Hacha showed a higher degree of differential gene regulation than Lachit when comparing excess Fe versus the control in both root and shoots. The difference between Hacha and Lachit was more pronounced in leaves than in roots (Supplemental Figure S2B). Among these genes slightly more genes were down-regulated than up-regulated (18.5 % and 15.6 % more in Hacha and Lachit leaves, respectively; 23.8 % and 0.5 % more in Hacha and Lachit roots, respectively), and more genes were differentially regulated in Hacha than in Lachit especially in leaves (26.4 % more in roots, 131.1 % more in leaves) (Figure 7A, excess Fe vs. control). To dissect the difference in response between the genotypes, differences between Hacha and Lachit were examined. In the comparisons between the two lines, 1147 and 1038 genes were more abundant in Lachit than in Hacha under the control condition and under excess Fe treatment, respectively. 1248 and 1161 genes were less abundant in Lachit than in Hacha roots under these conditions, respectively. Thus, under control conditions the number of differently abundant genes between Lachit and Hacha was slightly higher than under excess Fe treatment. In leaves, this difference was more pronounced. With a total of 939 genes expressed at higher and 890 genes expressed at lower level in Lachit than in Hacha under the control condition and 485 and 657 genes expressed lower in Lachit than in Hacha under these conditions, respectively. In summary, 37.6 % less genes were differently abundant between Lachit and Hacha under excess Fe in roots than under the control condition while this difference in roots was only 8.2 %. (Figure 7B, Lachit vs. Hacha). Thus, the numbers of genes, differently expressed between the two lines at a given Fe treatment, were relatively low, compared to the numbers of genes, regulated within the lines upon Fe stress versus control. A high number of genes was differentially expressed between excess Fe and the control condition within a line, but without being expressed at a different level between the two lines (e.g. 1462, 497 and 291 genes up in leaves, and respective other genes down in leaves, or up and down in roots; Supplemental Figure S3, Figure 7B). For this reason, we predict that these genes do not likely explain the major tolerance of Lachit in comparison with Hacha.

**Figure 7.**
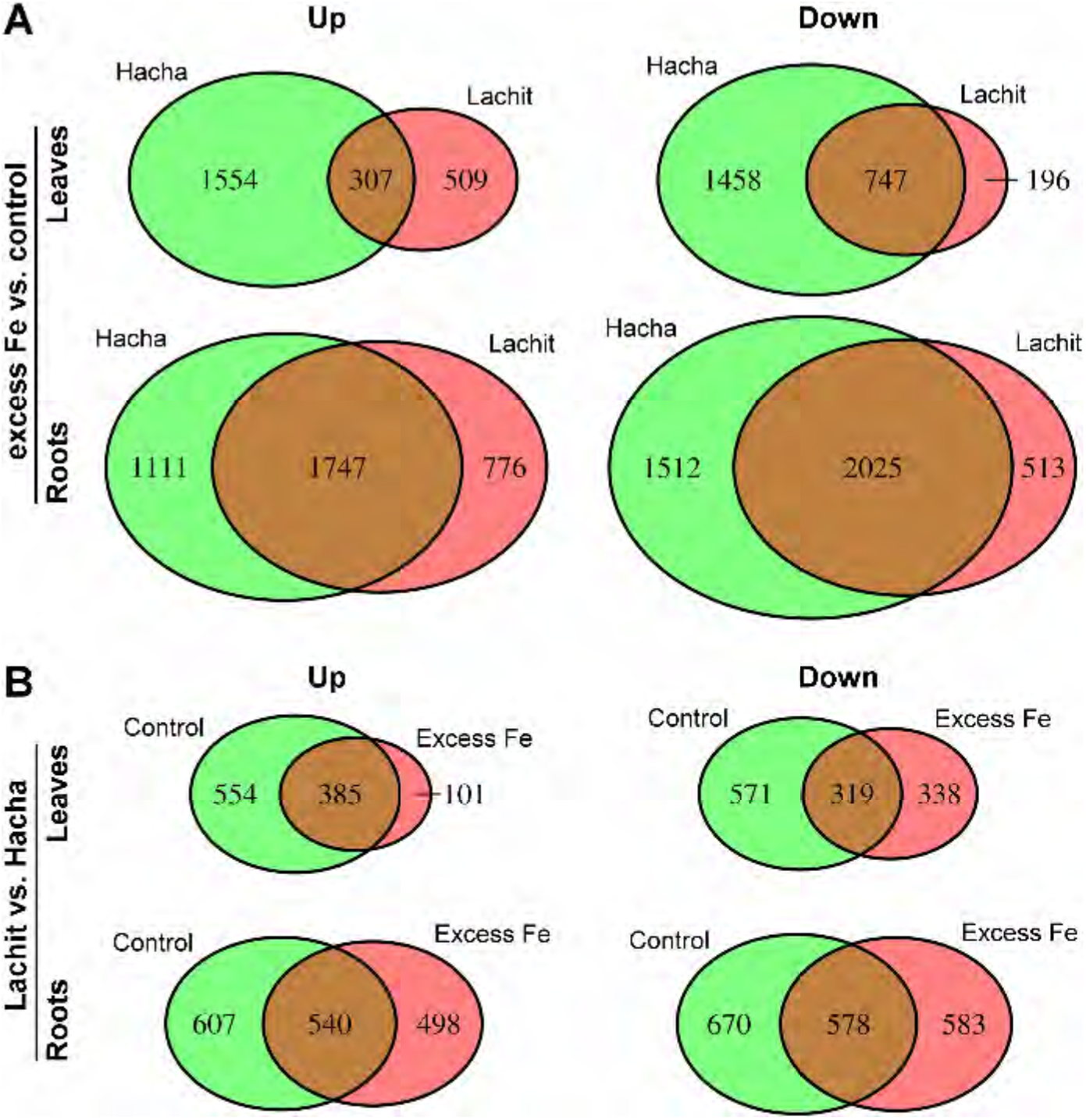
Two-way Venn diagrams with total numbers of regulated genes. (A), Excess Fe versus (vs.) control; (B), Lachit versus (vs.) Hacha. The areas of the diagram partitions reflect the numbers of up- and down-regulated (Up, Down) genes in the respective partitions.

We considered genes most interesting if they were differentially expressed between Hacha and Lachit. First of all, some of these gene groups are differentially expressed between excess Fe and control and at the same time differentially expressed between Lachit and Hacha (e.g. 92, 2 and 14 genes up in leaves of Hacha, Lachit or both in response to the stress and up in Lachit versus Hacha; similarly respective genes with corresponding downregulation, as well as genes showing these regulation patterns in roots; Supplemental Figure S3). A relatively large number of interesting genes was also differentially expressed in between Hacha and Lachit either under the control, excess Fe or both conditions, but without being regulated by the stress itself (e.g. 446, 94 and 380 genes up in leaves, and respective other genes down in leaves, or up and down in roots; Supplemental Figure S3). These differentially expressed genes that differentiate between Lachit and Hacha are most likely candidates for excess Fe tolerance of Lachit.

In summary, global transcriptome comparisons confirm that Hacha responded in a more drastic manner to excess Fe than Lachit. Leaf transcriptomes revealed more drastic responses to Fe stress in Hacha than in Lachit. However, the primary target of excess Fe in terms of total changes in gene expression patterns were roots rather than leaves. The transcriptome comparisons point to groups of genes, which may shed light on the tolerance mechanism in Lachit and explain the physiological phenotypes at the molecular level.

### Gene ontology (GO) term enrichment reflects morphological and physiological phenotypes of Hacha and Lachit in response to excess Fe

Multiple gene expression changes may be indicative of cellular pathways associated with variations in growth conditions or genotypes, commonly studied by gene ontology (GO) term enrichment.

At first, we investigated GO terms (Supplemental Tables S3-S14) related to plant performance. Since Hacha suffered more than Lachit during excess Fe treatment and ultimately died, we expected an enrichment of GO terms for cell breakdown in Hacha rather than in Lachit, in the comparisons of Fe excess stress versus the control. Indeed, we found an enrichment of 13 GO terms linked with cell death, organelle disassembly and autophagy in roots of Hacha (Figure 8A) and 16 such GO terms in leaves of Hacha (Figure 8B). In Lachit, 12 GO terms linked with cell death, organelle disassembly and autophagy were enriched in roots (Figure 8A) and none in leaves upon stress versus control (Figure 8B). In direct comparison of Lachit versus Hacha, six GO terms for cell death, organelle disassembly and autophagy were enriched in Hacha compared to Lachit in roots, but none in leaves (Figure 9). A cell death mechanism in plants is facilitated by reactive unsaturated fatty acid-derived lipid peroxide species, which are formed in the plasma membrane by lipoxygenases and in the presence of Fe (Dixon et al., 2012; Distéfano et al., 2017; Conrad et al., 2018). The group of 27 rice lipoxygenase genes (Supplemental Table S15) was significantly enriched among the 1861 genes up-regulated by excess Fe in Hacha leaves (Figure 7A; Fischer exact test, p = 0.01978) but not among the corresponding 816 genes in Lachit leaves, nor among any genes down-regulated by excess Fe in leaves nor in any of the root sample comparisons (Figure 7A). This confirms at the molecular level that Hacha leaves had been undergoing cell death in response to excess Fe in leaves.

**Figure 8.**
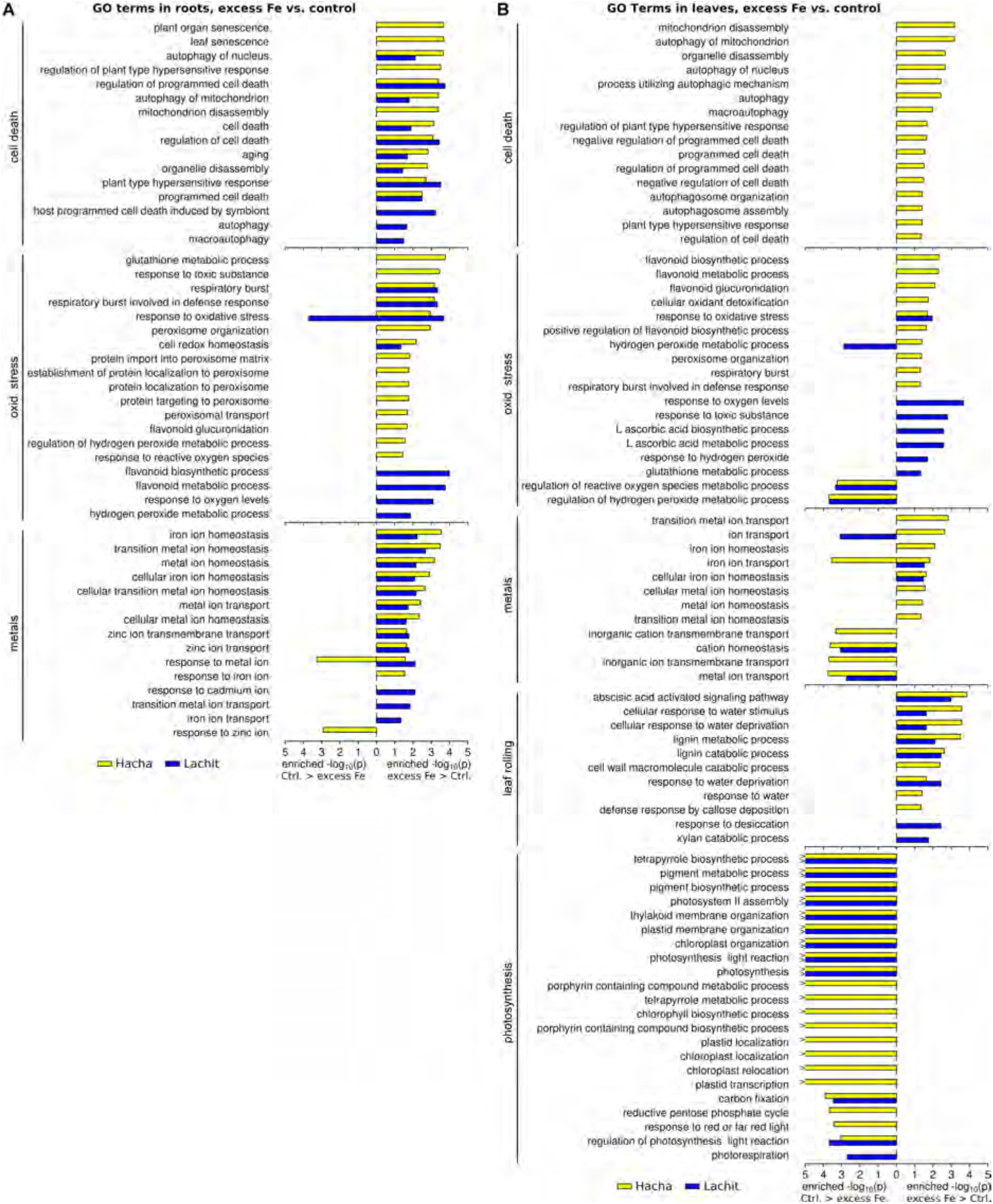
GO term enrichment of selected GO terms under excess Fe versus (vs.) the control. (A) Roots. (B) Leaves. Enrichment values are −log10(p) values; >, >1E-30 p value. GO term enrichment analysis was performed with R: topGO, see also Supplemental Tables S3 to S10.

**Figure 9.**
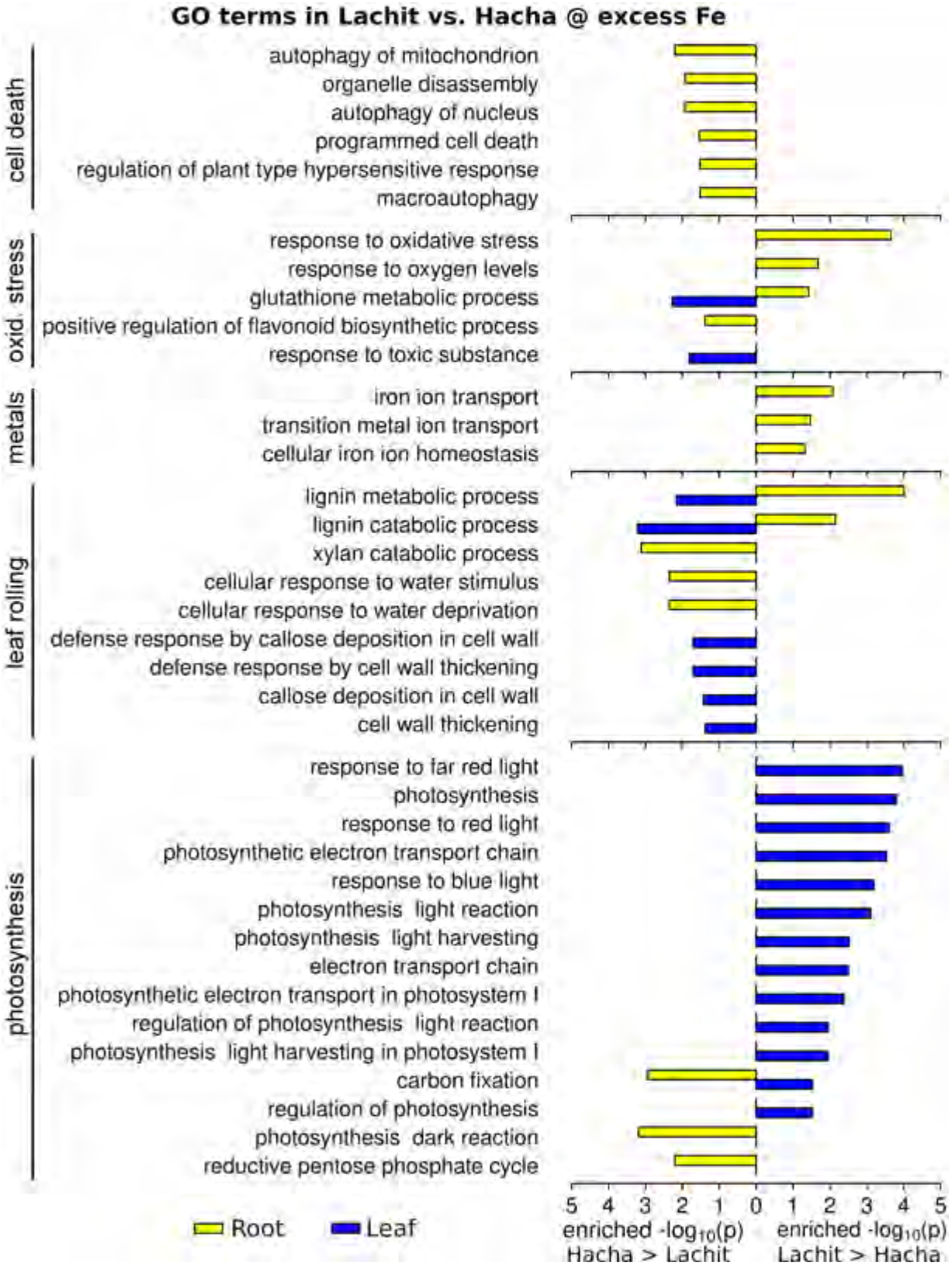
GO term enrichment of selected GO terms in Lachit versus (vs.) Hacha under excess Fe. Enrichment values are −log10(p) values; >, >1E-30 p value. GO term enrichment analysis was performed with R: topGO, see also Supplemental Tables S11 to S14.

Since Lachit in contrast to Hacha survived the excess Fe treatment, we expected an enrichment of GO terms for photosynthesis and light responses among the genes that were expressed at higher level in excess Fe-treated Lachit versus Hacha leaves. Indeed, 13 enriched GO terms were associated with photosynthesis and light responses in this comparison and, expectedly, none of them in roots (Figure 9). Thus, under excess Fe Lachit had continued to grow, as indicated by its responses to light and expressing photosynthesis functions compared to Hacha.

Hacha showed more signs of oxidative stress than Lachit in response to excess Fe. We therefore expected a molecular signature of oxidative stress in the comparison of Hacha and Lachit, in form of an enrichment of GO terms related to antioxidant metabolism, respiratory burst, redox homeostasis, peroxisomal functions, ROS responses and peroxidases. In Hacha roots and leaves, we found 15 and 10 enriched oxidative stress terms in response to Fe excess stress among up-regulated genes (Figure 8A, 8B), and two terms among down-regulated genes in leaves (Figure 8B). In the respective Lachit comparisons, there were 8 and 7 enriched terms among up-regulated genes, and three GO terms enriched among down-regulated genes (Figure 8A, 8B). Hence, Hacha enriched more oxidative stress-related GO terms than Lachit under excess Fe in roots and leaves. In the direct comparison between the lines, two oxidative stress terms were also enriched among genes that were up-regulated in Hacha versus Lachit leaves, and none was enriched in Lachit versus Hacha (Figure 9). However, in roots, three such terms were enriched among up-regulated genes in Lachit roots compared to Hacha roots under excess Fe, one term was down-regulated in roots (Figure 9).

Leaf rolling can be the result of sclerenchyma formation and drought (Cal et al., 2019). Hence, we expected lignin biosynthesis, cell wall, phenylpropanoid, abscisic acid and drought-related GO terms to be enriched in some of the comparisons. Indeed, in Fe excess-treated leaves versus control leaves 9 GO terms of these kinds were enriched in Hacha and 8 in Lachit leaves (Figure 8B). In the comparison of leaves under excess Fe treatment, six GO terms of these types were enriched among genes up-regulated in Hacha versus Lachit, and none among genes up-regulated in Lachit versus Hacha (Figure 9), whereby mostly cell wall thickening-related terms were enriched among genes expressed at a higher level in Hacha leaves compared to Lachit leaves (Figure 9). In the corresponding root comparisons, there were two GO terms related to lignin enriched among genes up-regulated in Lachit under stress, compared to three GO terms enriched among genes up-regulated in Hacha (Figure 9).

Leaf metal contents are the result of metal transport and allocation in the plant. The lower leaf metal contents of Lachit versus Hacha leaves in response to excess Fe may stem from differential metal homeostasis gene expression in roots and leaves. In Hacha roots, 11 metal homeostasis-related GO terms were enriched under excess Fe among up-regulated genes, two among down-regulated ones (Figure 8A), while in Lachit roots, there were 13 and zero GO terms enriched in respective comparisons (Figure 8A). In leaves, Hacha showed an enrichment of eight metal homeostasis-related GO terms among up-regulated genes, and five among down-regulated ones, whereas for Lachit only two such terms were enriched among up-regulated and three among down-regulated genes (Figure 8B). Hence, Hacha reacted more vividly to excess Fe than Lachit in leaves with respect to metal homeostasis-related GO terms. No metal homeostasis-related GO terms were enriched among differentially expressed genes in leaves, but three GO terms were enriched among genes up-regulated in Lachit versus Hacha in roots (Figure 9). This indicates that metal homeostasis genes expressed in roots might be involved in the Fe tolerance mechanism of Lachit.

Taken together, enrichment of different GO terms related to plant performance and cell death, oxidative stress, leaf rolling and drought as well as metal homeostasis reflect the different morphological and physiological behaviors of Hacha and Lachit in response to excess Fe.

### Individual target genes differentially expressed between Lachit and Hacha under excess Fe

To identify potential target genes for a metal tolerance mechanism we closely inspected genes from the oxidative stress (Supplemental Table S16) and metal homeostasis-related GO terms (Supplemental Table S17) that were enriched in Lachit compared to Hacha under excess Fe as up-regulated or down-regulated in this comparison. We also used several of these genes to validate the transcriptome set using reverse transcription (RT)-qPCR (Supplemental Figure S4).

Increased survival of Lachit may be caused by reduced root to shoot translocation, by decreased Fe uptake of the root, by increased tolerance of the stress, or by a combination of these. We examined potential Fe transport within the plant. In leaves, *YSL10* and Os07g0516600 were up-regulated in Lachit versus Hacha (2 genes in red circle in Figure 10 top left, Supplemental Table S18) while *HMA9, IPB1.2* as well as *PDR3* and *PDR5* were down-regulated (4 genes in red circle in Figure 10 top right, Supplemental Table S19). The regulation patterns of these genes suggest that they could be involved in keeping metal contents at a lower level in Lachit leaves than in Hacha leaves.

**Figure 10.**
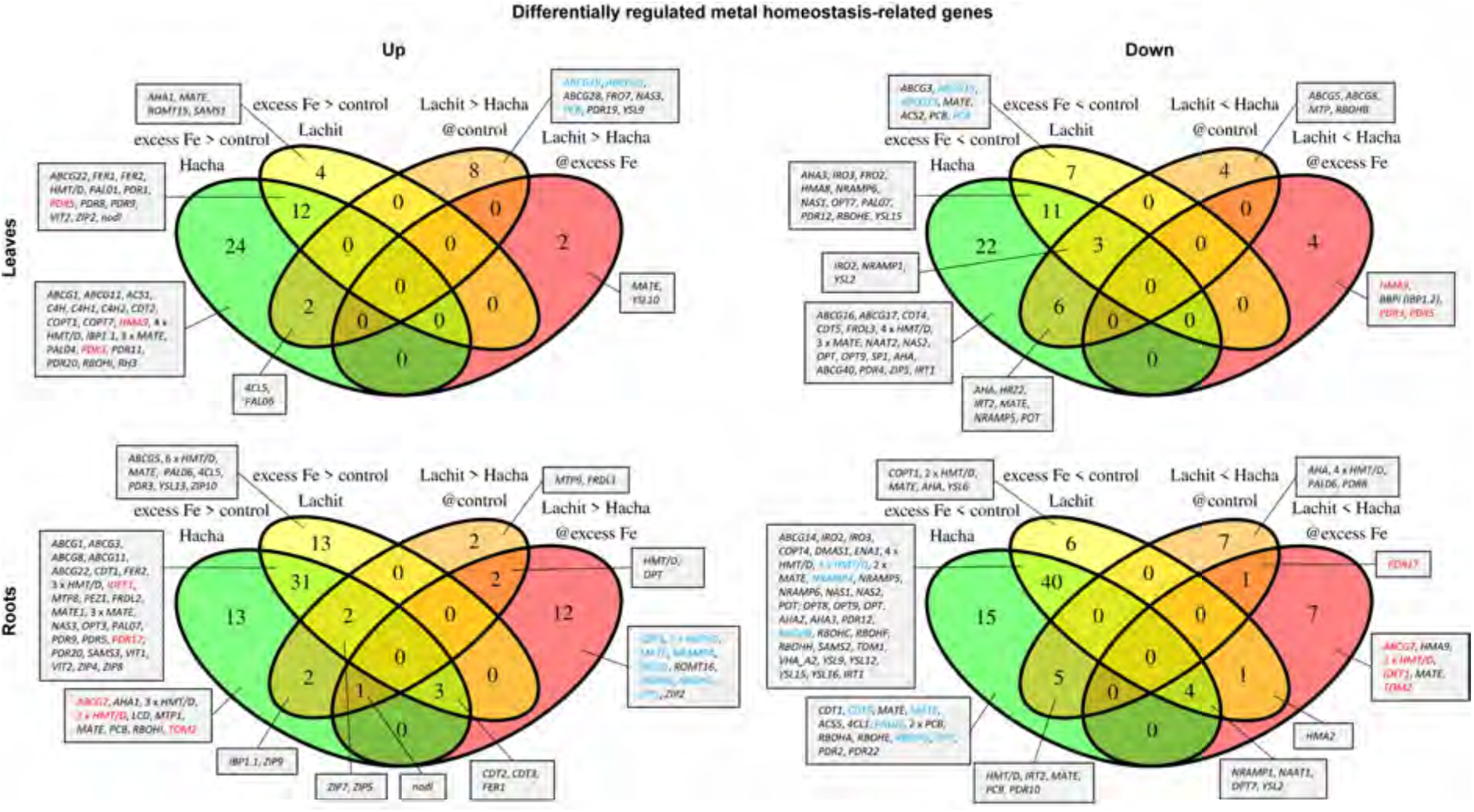
Four-way Venn diagrams of metal homeostasis-related genes. Left, up-regulated genes (Up); right, down-regulated genes (Down); upper part, leaves; lower part, roots. The respective comparisons are indicated. Metal homeostasis genes are indicated in grey boxes with gene name, gene family abbreviation or function abbreviation; blue gene names, expressed at higher level in Lachit than in Hacha (left) and down-regulated in Hacha (right); red gene names, expressed at lower level in Lachit than in Hacha (right) and up-regulated in Hacha (left). Arbitrary family or function abbreviations used above are PCB = phytochelatin biosynthesis, HMT/D = heavy metal transport/detoxification, nodl = nodulin-like. For detailed information on all the genes in this figure see Supplemental Tables S17-S21.

In excess Fe-stressed roots, 17 metal homeostasis genes were expressed at a higher level in Lachit than in Hacha, including four *HMT/D* genes, an *OPT* gene, a *MATE* transporter gene, *RBOHG, RBOHB, PAL02, ROMT16, NRAMP4, ZIP1, ZIP2, FER1*, OsNodulin-like2, *CDT2, CDT3* and *CDT5* (2, 12 and 3 genes in red circle in Figure 10 bottom left, Supplemental Table S20), while 13 genes were expressed at a lower level, like *PDR17, ABCG7*, two *HMT/D* genes, *MATE2, YLS2, OPT7, NRAMP1, NAAT1, HMA2, HMA9* and *TOM2* (1, 7, 1 and 4 genes in Figure 10 bottom right, Supplemental Table S21). Therefore, the regulation patterns of these genes suggest that they might contribute to exclusion of metals from root and from root to shoot transport in Lachit rather than in Hacha.

Finally, we tested differences in the stress responses. We assembled a list of oxidative stress-related genes in Venn diagrams (Supplemental Figure S5, Supplemental Table S16, Supplemental Tables S22 to S25). Different members of gene families coding for isozymes involved in antioxidant reactions frequently were found expressed in opposite manner. For example, one *AO* (coding for ascorbate oxidase), one *PRX* (coding for a peroxidase) and one *DHAR* gene (encoding a dehydroascorbate reductase) expressed at a higher level in Lachit compared to Hacha leaves, whereas a *MDAR* (coding for a monodehydroascorbate reductase) stayed neutral, one *TRX-PRX* (coding for thioredoxin peroxidase) and six *GST* genes (encoding glutathione-S-transferases) were expressed at a lower level in Lachit compared to Hacha leaves. A clear regulation pattern of such gene functions was therefore not apparent.

Taken together, close inspection of differential expression of metal homeostasis- and oxidative stress-related genes strongly indicates a tolerance mechanism that is based rather on diminished metal uptake and retention of Fe and other bivalent metals in the root tissues and enhanced local Fe and metal cellular detoxification rather than a tolerance mechanism based on antioxidant genes. A root-borne metal homeostasis mechanism may be in place because of the higher number of differentially expressed genes related to this category in Lachit versus Hacha. The identified 6 and 30 metal homeostasis genes expressed differentially between Lachit and Hacha in leaves or roots (total of 35 different genes; Table 1) were thus targets for metal tolerance in Lachit.

**Table 1.** SNP analysis of 35 metal homeostasis-related and putative iron tolerance genes with regard to selection during rice domestication and breeding

| Gene | MSU ID | RAP ID | 1 | 2 | 3 | 4 |
| --- | --- | --- | --- | --- | --- | --- |
| *IBP1* | Os01g0124401 | LOC_Os01g03360 |  | YES |  |  |
| *HMT/D protein* | Os01g0125600 | LOC_Os01g03490 | YES |  | YES |  |
| *CDT3* | Os01g0178300 | LOC_Os01g08300 |  |  |  | YES |
| *RBOHB* | Os01g0360200 | LOC_Os01g25820 |  |  |  | YES |
| *ZIP1* | Os01g0972200 | LOC_Os01g74110 |  |  |  |  |
| *NRAMP4* | Os02g0131800 | LOC_Os02g03900 | YES |  | YES | YES |
| *NAAT1* | Os02g0306401 | LOC_Os02g20360 |  |  |  | YES |
| *YSL2* | Os02g0649900 | LOC_Os02g43370 |  | YES |  | YES |
| *ZIP2* | Os03g0411800 | LOC_Os03g29850 |  |  |  | YES |
| *OPT7* | Os03g0751100 | LOC_Os03g54000 |  |  |  |  |
| *HMT/D protein* | Os03g0819400 | LOC_Os03g60480 |  |  |  | YES |
| *MatE protein* | Os03g0839200 | LOC_Os03g62270 |  |  |  |  |
| *ABCG7* | Os03g0859500 | LOC_Os03g64200 |  |  |  |  |
| *PAL2* | Os04g0518100 | LOC_Os04g43760 |  |  |  |  |
| *YSL10* | Os04g0674600 | LOC_Os04g57840 | YES |  | YES | YES |
| *OsNodulin-like2* | Os04g0686800 | LOC_Os04g59020 | YES |  |  | YES |
| *CDT5* | Os05g0178300 | LOC_Os05g08554 | YES |  |  | YES |
| *MATE2* | Os05g0554000 | LOC_Os05g48040 |  |  |  |  |
| *CDT2* | Os06g0143100 | LOC_Os06g05120 |  |  |  |  |
| *ROMT16* | Os06g0165800 | LOC_Os06g06980 |  |  |  |  |
| *OPT protein* | Os06g0239300 | LOC_Os06g13200 |  |  |  | YES |
| *HMA9* | Os06g0665800 | LOC_Os06g45500 | YES |  | YES | YES |
| *HMA2* | Os06g0700700 | LOC_Os06g48720 |  |  |  |  |
| *NRAMP1* | Os07g0258400 | LOC_Os07g15460 |  |  |  |  |
| *MatE protein* | Os07g0516600 | LOC_Os07g33310 |  |  |  | YES |
| *PDR5* | Os07g0522500 | LOC_Os07g33780 |  |  |  |  |
| *HMT/D protein* | Os07g0682000 | LOC_Os07g48390 |  |  |  | YES |
| *IDEF1* | Os08g0101000 | LOC_Os08g01090 |  |  |  | YES |
| *HMT/D protein* | Os08g0403300 | LOC_Os08g31140 |  |  |  | YES |
| *HMT/D protein* | Os09g0272000 | LOC_Os09g09930 |  |  |  | YES |
| *RBOHG* | Os09g0438000 | LOC_Os09g26660 | YES |  |  | YES |
| *FER1* | Os11g0106700 | LOC_Os11g01530 |  |  |  |  |
| *TOM2* | Os11g0135000 | LOC_Os11g04030 |  |  |  | YES |
| *PDR3* | Os11g0587600 | LOC_Os11g37700 |  |  |  |  |
| *HMT/D protein* | Os12g0144600 | LOC_Os12g05040 | YES |  |  | YES |

### Enrichment of metal homeostasis candidate genes during rice domestication

Beneficial genes for agronomic purposes are likely targets during domestication and breeding. To obtain functional hints about the aforementioned 35 gene functions reflecting metal tolerance in Lachit (Table 1), we examined their presence among genes which had been selected during breeding and speciation of *O. rufipogon* and *O. sativa*. Interestingly, we found that eight genes were included in the selective sweep regions according to (Huang et al., 2012), namely *Os01g0125600* (Heavy metal transport/detoxification protein domain containing protein), *NRAMP4, YSL10, OsNodulin-like2* (similar to *AtVIT* homolog 4), *CDT5, HMA9, RBOHG* and *Os12g0144600* (Heavy metal transport/detoxification protein domain containing protein) (Table 1, Supplemental Table S26). Furthermore, two genes showed breeding signatures of rice improvement according to (Xie et al., 2015), *IBP1* and *YSL2* (Table 1, Supplemental Table S26). We separately performed SNP haplotype analysis of 147 rice accessions against *O. rufipogon* and found that four genes were included in selective sweep regions in the speciation of *O. rufipogon* and *O. sativa, Os01g0125600* (Heavy metal transport/detoxification protein domain containing protein), *NRAMP4, YSL10* and *HMA9* (Table 1, Supplemental Table S26).

Since we investigated Indica varieties and our results differed from previous studies in which Japonica varieties had been investigated (L. B. Wu, Ueda, Lai, & Frei, 2017), we asked whether the Fe tolerance genes found in this study were subject to differentiation between Indica and Japonica rice varieties. Phylogenetic SNP analysis of the 35 genes was performed, which showed that 20 of them were subject to differentiation between Indica and Japonica varieties. These genes include heavy metal transport/detoxification protein domain-containing proteins (*Os03g0819400, Os07g0682000, Os08g0403300, Os09g0272000* and *Os12g0144600), CDT3, CDT5, RBOHB, NRAMP4, NAAT1, YSL2, ZIP2, YSL10, OsNodulin-like2* (similar to *AtVIT homolog 4*), an Oligopeptide transporter (*OPT*) domain-containing gene (*Os06g0239300), HMA9*, a Multi antimicrobial extrusion protein *MatE* gene (*Os07g0516600), IDEF1* and *RBOHG* (Table 1, Supplemental File 1).

Taken together, 22 different genes out of the 35 metal homeostasis candidates had been subject of selection during domestication and breeding.

## Discussion

Here, we report a tolerance mechanism of an Indica rice variety to high Fe stress related to root-borne metal homeostasis. We selected and characterized a pair of Indica rice varieties with contrasting abilities to cope with Fe excess stress. Hacha was more susceptible than Lachit, while Lachit, although it was affected, was able to tolerate the Fe excess stress. High Fe tolerance genes were selected during rice domestication.

In both, Hacha and Lachit, Fe excess caused reduction in leaf growth and leaf rolling phenotypes. Excessive amounts of Fe were transported to leaves where Fe accumulated to high levels, particularly in the vascular bundles and in sclerenchyma tissue. Excess Fe acted as a redox-reactive ion and caused H_2_O_2_ generation and lipid peroxidation. More than 7.000 genes were differentially expressed in roots and leaves of Fe excess-treated plants versus control plants, showing the profound effect of this stress on plant growth and physiology also at the molecular level. These molecular patterns explain the physiological phenotypes. Oxidative stress and Fe transport GO terms were induced in the Fe excess stress-treated plants, showing that plants had perceived and responded to Fe and ROS signals. Leaf rolling could be related to dehydration stress, as regulated GO terms for ABA and drought responses were apparent. On the other hand, cell wall modification and lignification may also be relevant for leaf rolling, and similarly resulted in a Fe excess-dependent changes of GO terms for phenylpropanoid metabolism and cell wall modification (Cal et al. 2019). Plant performance was severely affected and the lower growth and compromised survival under Fe excess correlated with lower photosynthesis-related gene expression and cell death-related GO term regulation. These symptoms were expected because of the cytotoxic redox effects of elevated Fe in cells and plants, reported before (Tewari, Hadacek, Sassmann, & Lang, 2013).

A very interesting question was to explore the differences between Hacha and Lachit and the mechanisms behind the tolerance of Lachit to excess Fe stress. We noted that with regard to all Fe stress symptoms Hacha had responded in a more severe manner than Lachit to the stress (Table 2). We interpret the findings as follows: Hacha transports more Fe, Zn, Mn and Cu from root to leaves than Lachit under Fe excess stress. Consequently, leaves respond in a stronger manner by leaf rolling and lipid peroxidation. The toxic effects of Fe are stronger in Hacha than in Lachit, ultimately leading to plant growth failure of Hacha but not of Lachit. These morphological and physiological outcomes are triggered by gene expression changes. The total number of differentially regulated genes in leaves and roots was about 50 % more elevated in Hacha compared with Lachit. These genes, being differentially expressed in Hacha versus Lachit upon the stress, reflected several GO terms for poor plant performance, such as cell disassembly and cell death, lipoxygenase, drought and oxidative stress. On the other side, the better growth of Lachit was mirrored by an enrichment of GO categories related to photosynthesis.

**Table 2.**
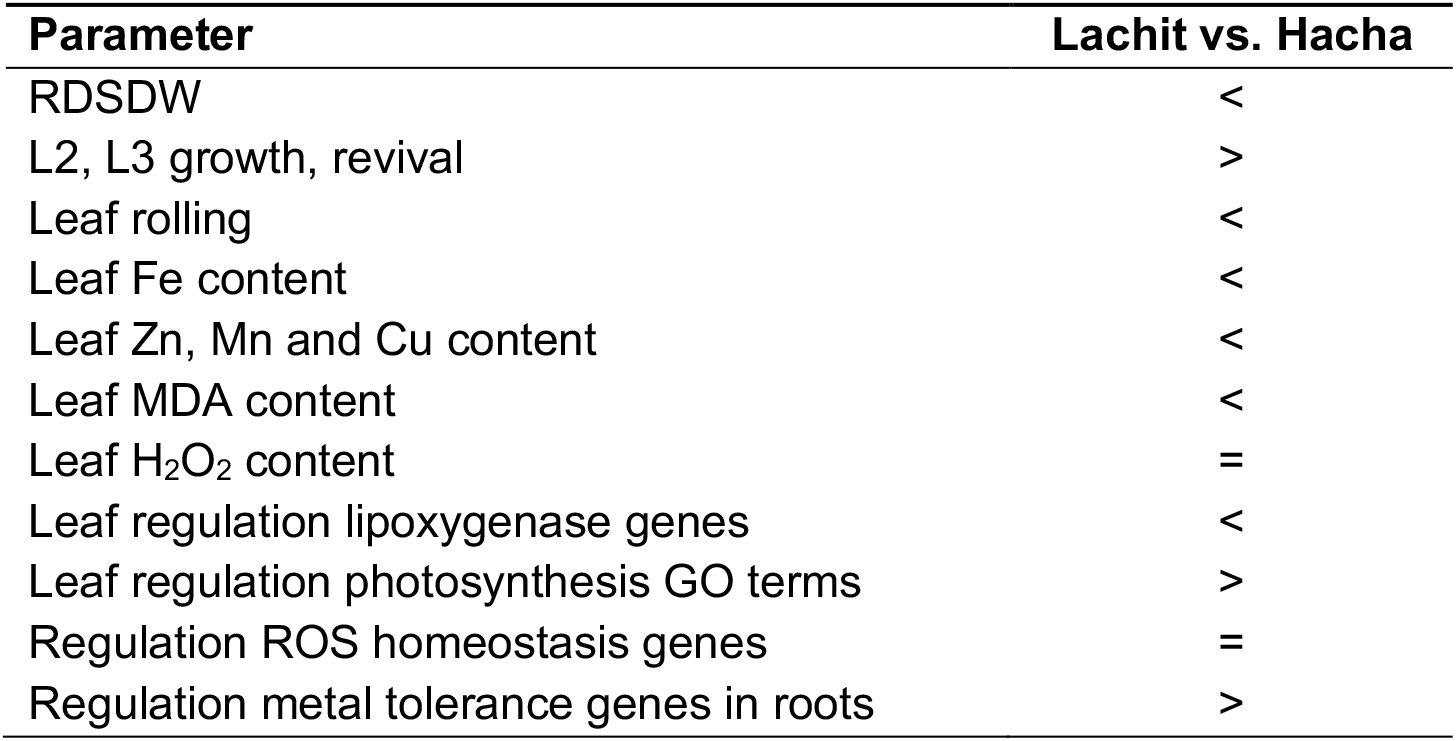
Summary of responses of Hacha and Lachit to excess Fe

When scanning regulated genes between Lachit and Hacha under excess Fe, we could detect differences in 35 metal homeostasis genes that may explain tolerance. Several genes are indicators for a mechanism that relates to heavy metal detoxification in root cells. Ferritin binds and detoxifies Fe (Briat, Duc, Ravet, & Gaymard, 2010), and *FER1* was found up-regulated in Lachit versus Hacha roots, indicating Fe retention, Fe detoxification in the root tissues and protection from oxidative stress. Moreover, three out of four rice *CDT* genes are expressed at higher levels in Lachit versus Hacha roots under excess Fe. *CDT3* might be regulated by ART1, an Al tolerance transcription factor (Xia, Yamaji, & Ma, 2013). In our experiment, *ART1* was not regulated in Hacha or Lachit roots. Therefore, we conclude that induction of *CDT3* was caused by other factors. The regulation pattern of *CDT2* and *CDT3* imply that they might be co-regulated in an ART1-independent manner, whereas *CDT5* was clearly not co-regulated with *CDT2* or *CDT3* under excess Fe. Although the exact function of these proteins is not yet known, CDTs may bind metal ions due to a cysteine-rich C-terminal domain. OsCDT2.2 may chelate Cd in the apoplast, impeding Cd uptake into the cell (Kuramata et al., 2008). OsCDT3 has Al-binding activity, which may occur at the inner side of the plasma membrane (Xia et al., 2013). Concomitantly, CDT proteins may sequester Fe in the apoplast or in the cell. Furthermore, *OsNodulin-like2*, an ortholog of *AtVTL4*, was expressed at a higher level in Lachit than in Hacha roots. Being a putative vacuolar Fe transporter, *OsNodulin-like2* could contribute to detoxification and retention of Fe in the root tissues by increased sequestration of Fe in the vacuole. Taken together, *CDT*, *FER1* and *OsNodulin-like2* genes may all contribute to Fe tolerance as well as tolerance to other heavy metals and Al (Kuramata et al., 2008; Xia et al., 2013). *CDT* and *OsNodulin-like2* are therefore excellent candidate genes for further research on Fe toxicity and tolerance in rice and other plant species. This is particularly interesting since Fe and Al toxicity both occur in lowland rice grown in acidic soils, suggesting that Fe and Al tolerance are linked. Other genes out of the 35 metal homeostasis genes point to a second mechanism for excess Fe tolerance in Lachit which is the exclusion of Fe, Mn, Zn and Cu from xylem-mediated transport to shoots. Decreased Zn, Mn and Cu contents in response to high Fe were reported before (Shao, Chen, Wang, Mou, & Zhang, 2007; Tanaka et al., 1966). But it is unexpected that this metal interference effect with retention of metals in the root occurred in Lachit but much less in Hacha. Indeed, roots present the first defense line to excess metals. YSL2 nicotianamine transporter plays a role in root-to-shoot translocation of Fe (Ishimaru et al., 2010). Phytosiderophore efflux transporter TOM2 is also important for Fe distribution from the root to the shoot (Nozoye et al., 2015). *YSL2* and *TOM2* genes were expressed at lower levels in Lachit roots. Several genes for Fe acquisition from the soil into the root were expressed at lower level in Lachit than in Hacha roots under excess Fe, suggesting that Lachit may have taken up less Fe from the environment than Hacha in the excess Fe condition. These genes encode enzymes for phytosiderophore synthesis, e.g. nicotianamine amino transferase (*NAAT1*) (T. Kobayashi, Nakanishi, Takahashi, Mori, & Nishizawa, 2008). Down-regulation of phytosiderophore biosynthesis would also explain lower contents of Cu and Zn (Nozoye et al., 2015; Widodo et al., 2010). Some metal transporter genes, *NRAMP1, HMA2* and *HMA9* were also expressed at lower level in Lachit than in Hacha roots. NRAMP1 has been shown to act in cellular Cd uptake and distribution from the root to the leaves (Takahashi et al., 2011). *NRAMP1* was previously found induced under Fe deficiency, and hence, may play a role in Fe redistribution to shoots (Bashir et al., 2014). Oligopeptide transporter *opt7* knockout plants accumulate more Fe in the shoot than wild type, and Fe is not available under Fe-replete conditions (Bashir et al., 2015). HMA2 participates in root-to-shoot transport of Zn and Cd (Takahashi, Bashir, Ishimaru, Nishizawa, & Nakanishi, 2012; Takahashi, Ishimaru, et al., 2012). Some questions on the function of HMA2 and OPT7 need to be solved, since their lower expression should lead to higher Fe, Cu and Zn in leaves, contradicting metal contents found in Lachit. PDR17, ABCG7, the two HMT/D proteins and multidrug and toxic compound extrusion transporter, MATE2, have not been characterized in the context of metal homeostasis. Among the other genes differentially regulated in roots in Lachit versus Hacha, *RBOHG, RBOHB* and the *HMT/D*, other *OPT* and *MATE* genes have not been sufficiently characterized in the context of metal homeostasis in rice. NADPH oxidase/respiratory burst oxidase homolog (RBOH) proteins generate localized ROS increases to regulate developmental and stress responses (Chapman, Muhlemann, Gayomba, & Muday, 2019), and down-regulation upon high Fe may prevent excessive ROS signaling. Phenylalanine ammonia lyase PAL02 and caffeoyl-CoA-O-methyltransferase ROMT16 (CCoAOMT-1) act in concert in the coumarin biosynthesis pathway. Coumarins are known to play a role in Fe uptake in Strategy I plants (Bashir et al., 2011; Fourcroy et al., 2014; Ishimaru, Bashir, Nakanishi, & Nishizawa, 2011; Ishimaru, Kakei, et al., 2011; Rajniak et al., 2018; Siso-Terraza et al., 2016; Tsai et al., 2018; Ziegler, Schmidt, Strehmel, Scheel, & Abel, 2017). Their role in Strategy II plants in Fe homeostasis in unclear. NRAMP4 is an aluminum uptake transporter (Xia, Yamaji, Kasai, & Ma, 2010). Metal transporter ZIP2 has not been functionally characterized itself, but its homolog ZIP1 has been shown to transport different metal ions (Ramesh, Shin, Eide, & Schachtman, 2003) and has later been found to act as a plasma membrane and ER-located Zn, Cu and Cd efflux transporter (Liu et al., 2019). Although ZIP1 probably does not transport Fe (Liu et al., 2019), higher expression of *ZIP1* and *ZIP2* in the tolerant versus the sensitive line under excess Fe suggests that they might contribute to the tolerance mechanism.

A really interesting question is how these metal homeostasis genes are regulated in the root in response to excess Fe. As explained above, IDEF1 up-regulates Fe acquisition genes under low Fe (Takanori Kobayashi et al., 2009; Takanori Kobayashi et al., 2007). Interestingly, *YSL2* is one of the downstream genes of IDEF1. Hence, repression of *IDEF1* might have contributed to lower expression of *YSL2* and consecutively to reduced root-to-shoot allocation of Fe. Obviously, some metal homeostasis genes are regulated similarly in Hacha and Lachit while only a subset of the genes is differentially regulated and thus expressed for example at lower level by excess Fe in Lachit than in Hacha. Since the transcription factor IDEF1 is among them, IDEF1 represents a candidate as a target for heavy metal tolerance regulation.

Among the leaf-regulated genes Os07g0516600 protein, a multi-antimicrobial extrusion (MatE) domain-containing protein, YSL10, a YELLOWSTRIPE-LIKE putative nicotianamine-metal chelate transporter, transporters PDR3 and PDR5, belonging to the family of ATP-binding cassette transporter and pleiotropic drug resistance, have not been characterized yet. *HMA9* is induced by high Zn, Cu and cadmium (Cd), and the encoded heavy metal ATPase protein, may serve as an efflux transporter for metal detoxification (Lee, Kim, Lee, & An, 2007). IBP1.2 and its close homolog and trypsin inhibitor IBP1.1 interact with the transcription factor IDEF1, up-regulating Fe uptake in rice roots in response to -Fe (Takanori Kobayashi et al., 2009; Takanori Kobayashi et al., 2007). IBP1.1 prevents 26S proteasome-dependent degradation of IDEF1 (L. Zhang et al., 2014). The role of IBP1.1 and IBP1.2 in leaves is not elucidated.

In a previous study, tolerance to an overload in Fe was found associated with an increase of enzymatic antioxidants, like catalase, glutathione reductase and superoxide dismutase, and non-enzymatic compounds, such as ascorbic acid, glutathione and phenolic compounds, (L. B. Wu et al., 2017). This tolerance mechanism was apparent at the genetic level displayed by differential gene regulation between tolerant and sensitive lines. However, in our study, the genes encoding isozyme components for the ascorbate pool showed varying and non-conclusive expression patterns. One *DHAR* and *MDHAR* gene were expressed at higher level in Lachit than Hacha leaves under Fe stress, while other genes of this kind were oppositely regulated, and neither *DHAR* nor *MDHAR* was regulated in roots, similar for *AO* genes. The antioxidant system may therefore not be the primary mechanism for Fe stress tolerance in Lachit which would speak against a suggested shoot-borne Fe tolerance mechanism related to antioxidant enzymes in Lachit (L. B. Wu et al., 2017).

Albeit indirectly, selection sweep region, breeding signature and SNP haplotype analyses suggest a certain importance of 22 out of the 35 potential Fe tolerance genes. These observations indicate functional significance of metal homeostasis genes identified in our study due to the correlation with lowland rice breeding. Beneficial traits are usually selected during speciation of wild and cultivated rice and breeding of agriculturally important varieties. Additionally, analysis of phylogenetic tree differentiation revealed that 20 out of the 35 Fe tolerance genes are substantially different in Indica and Japonica rice accessions. This may suggest that metal homeostasis genes for Fe excess tolerance may play an important role in rice differentiation and breeding.

In summary, our study highlights new candidate genes for Fe excess tolerance in the lowland rice variety Lachit. These include genes that have not been associated with Fe homeostasis before. The most noteworthy gene functions standing out from our data comprise reduced mobilization as well as increased retention of Fe and other bivalent metals in the root tissues. As this was confirmed by reduced shoot metal contents and GO term enrichment analysis, we conclude that the main Fe tolerance mechanism in Lachit is based on these functions. The respective genes can be explored as targets for breeding of improved lowland rice crops. In the perspectives of this work, the importance of individual genes can be further confirmed in transgenic and genetic experiments. Likewise, it would also be interesting to determine why and how these genes are regulated differently in the cultivars.

## Acknowledgements

The authors thank Elke Wieneke for technical assistance. This work received funding from Germany’s Excellence Strategy, EXC 2048/1, Project iD: 390686111. The authors have no conflict of interest to declare.

## Author contributions

H.-J.M., A.B., S.K.P., P.B. conceived the project; S.K., H.-J.M., H.K., H.B.A., S.F., C.F.-S. performed experiments; A.B., H.-J.M., S.K., H.B.A., A.B., S.K.P., G.X., L.S. and P.B. analyzed the data; P.B. and H.-J.M. wrote the original draft; and H.-J.M., A.B., S.K.P., G.X., L.S. and P.B. reviewed and edited the article; P.B. acquired funding.

## Supplemental Figures

**Supplemental Figure S1.**
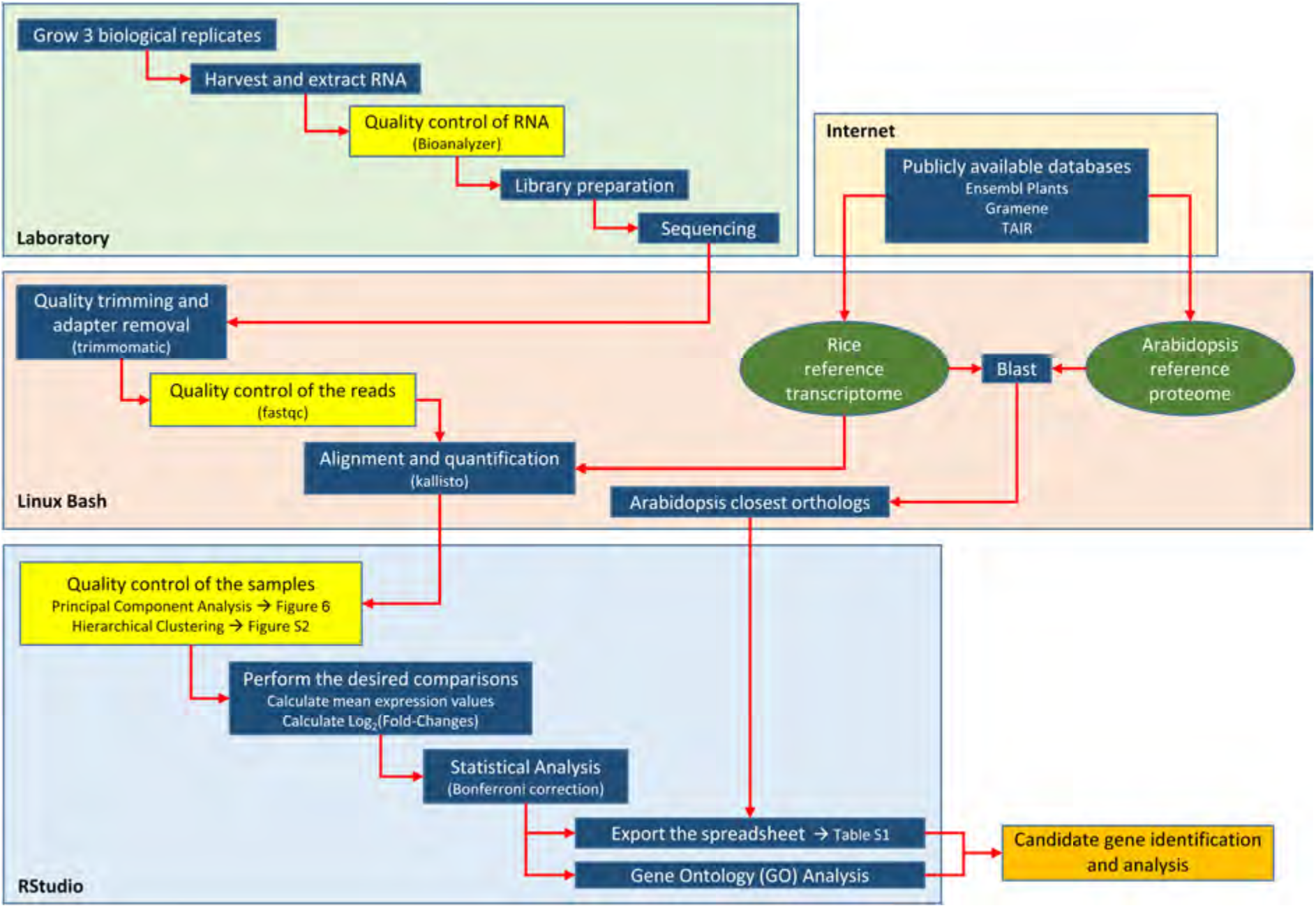
Work flow of RNA-Seq experiment. Three biological replicates of eight Hacha and Lachit rice plants each were grown. After the Fe excess and control treatment period, the plants were harvested and RNA was extracted. The quality of the RNA was checked with a Bioanalyzer. After library preparation and sequencing the output was trimmed by quality and adapter sequences were removed. The quality of the reads was checked using fastqc. After preparing the kallisto index with the rice reference transcriptome, the reads were aligned and quantified in paired-end mode using kallisto. The resulting TSV files containing the transcript identifiers and expression TPM (transcripts per million) values were read into R using RStudio. The quality of the samples was checked by PCA and HC, and data statistically evaluated. The closest Arabidopsis orthologs were determined by blasting each rice transcript against the Arabidopsis reference proteome and added to the existing data. Finally, Table S1 was exported as a spreadsheet file containing all the original and processed data and information. GO term enrichment analyses were conducted, and further analyses performed in and outside R.

**Supplemental Figure S2.**
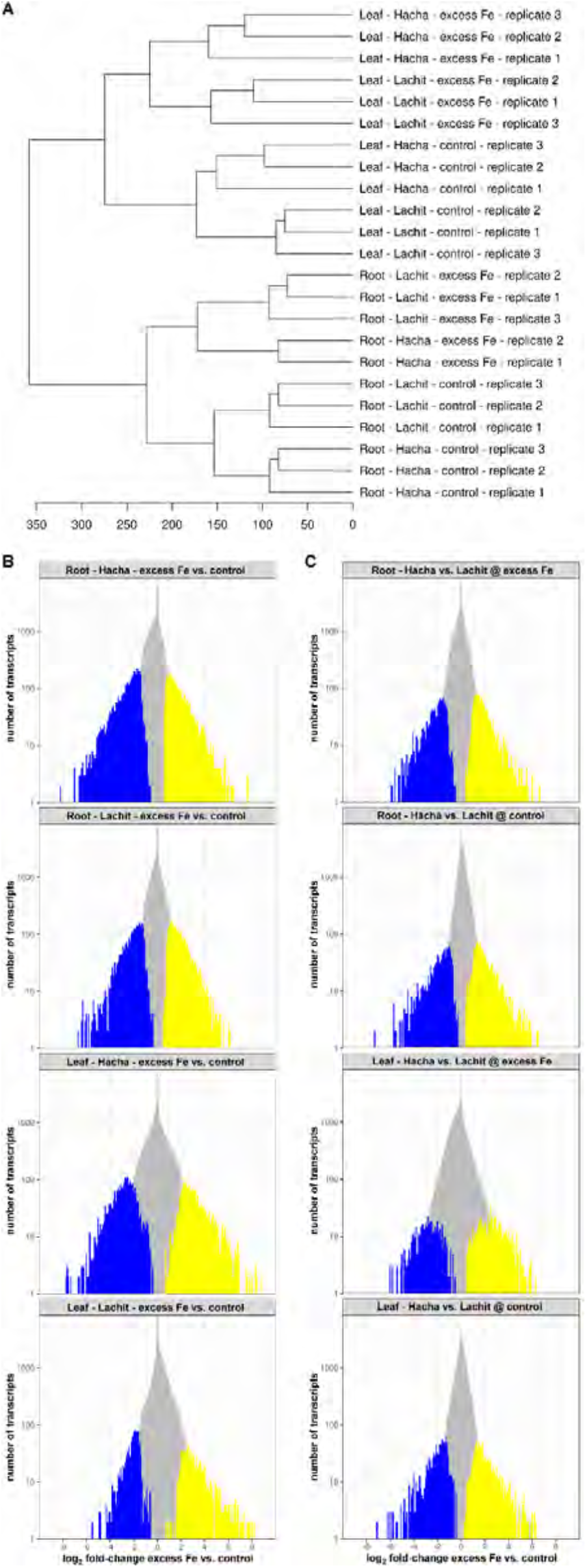
Quality control of the samples and overview of the numbers of regulated transcripts. (A) Dendrogram of hierarchical clustering (HC) of the samples. (B) Histograms of regulated transcripts with log_2_(fold-change) values in excess Fe vs. control comparisons in roots and leaves. (C) Histograms of regulated transcripts with log_2_(fold-changes) in comparisons between Hacha and Lachit in roots and leaves at excess Fe and control conditions, respectively. (B-C) Yellow bars represent significantly up-regulated transcripts and blue bars represent significantly down-regulated transcripts (p < 0.01).

**Supplemental Figure S3.**
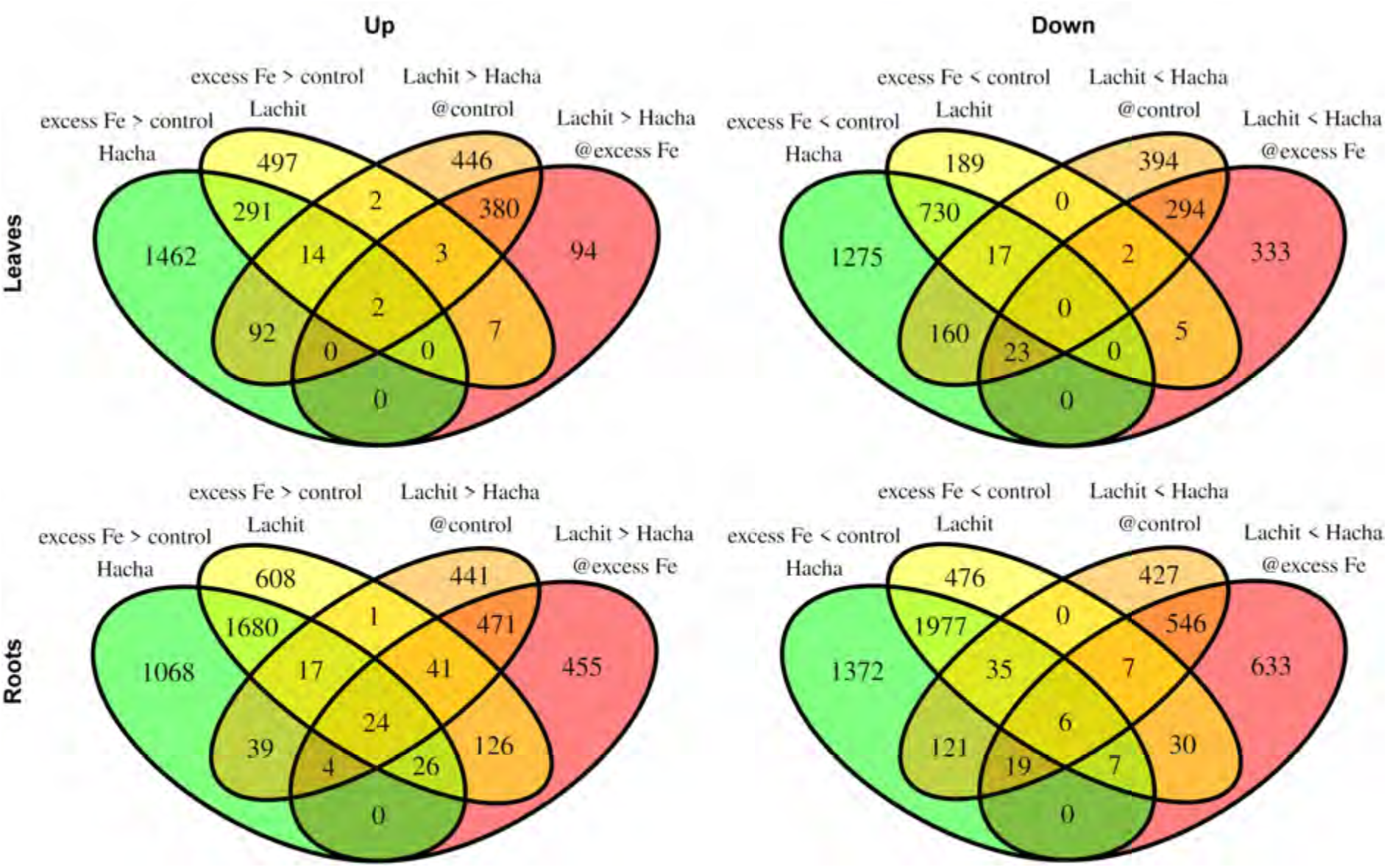
Four-way Venn diagrams with all meaningful comparisons performed. Left, up-regulated genes (Up); right, down-regulated genes (Down); upper part, leaves; lower part, roots. The respective comparisons are indicated. The numbers of genes with respective expression patterns are noted.

**Supplemental Figure S4.**
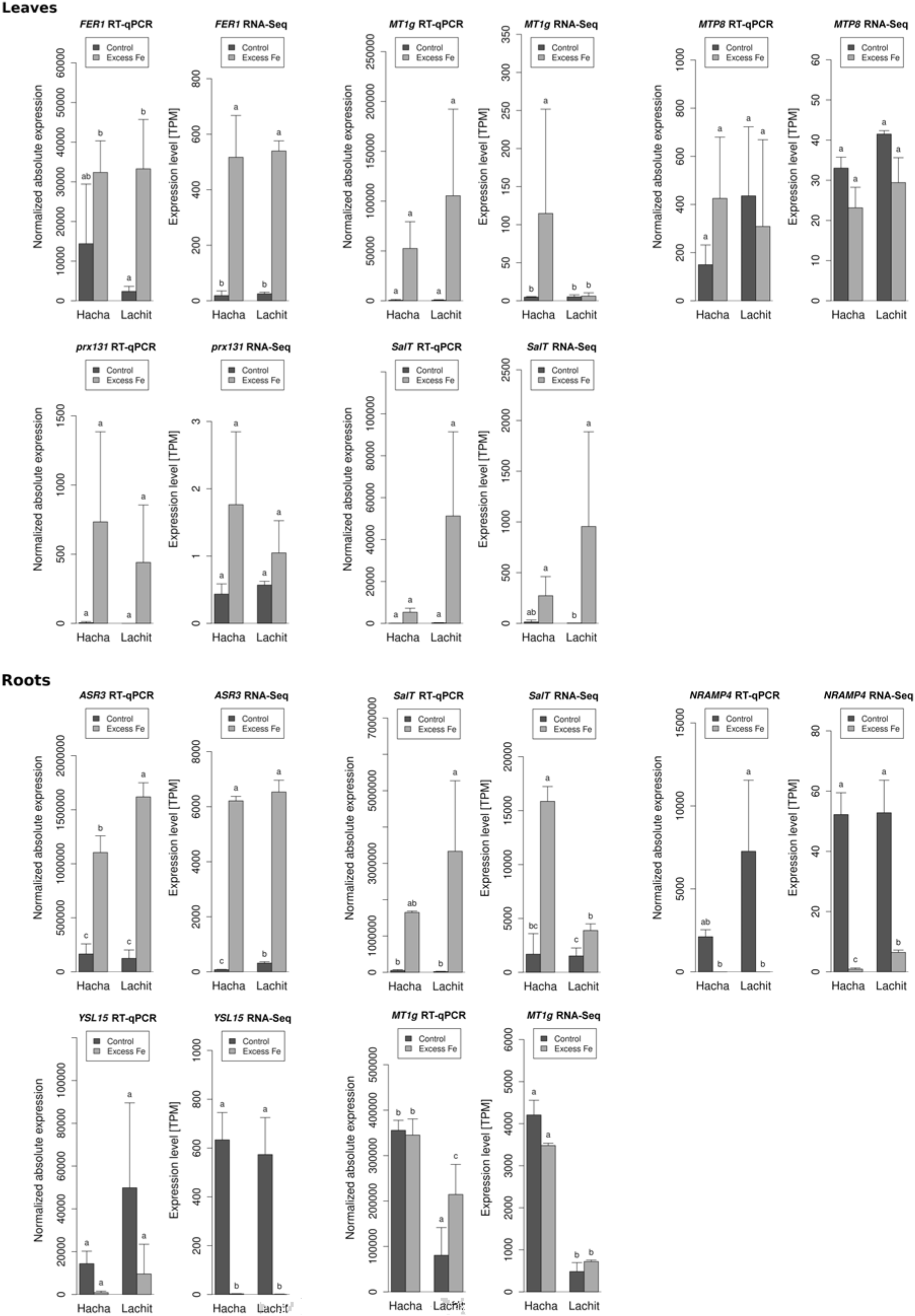
Validation of RNA-seq data by reverse transcription (RT)-qPCR. Gene expression of selected genes was investigated and compared for RT-qPCR (left) and RNA-seq (right). Different letters indicate a significant difference in normalized absolute expression levels (p < 0.05 in RT-qPCR and q < 0.01 in RNA-Seq).

**Supplemental Figure S5.**
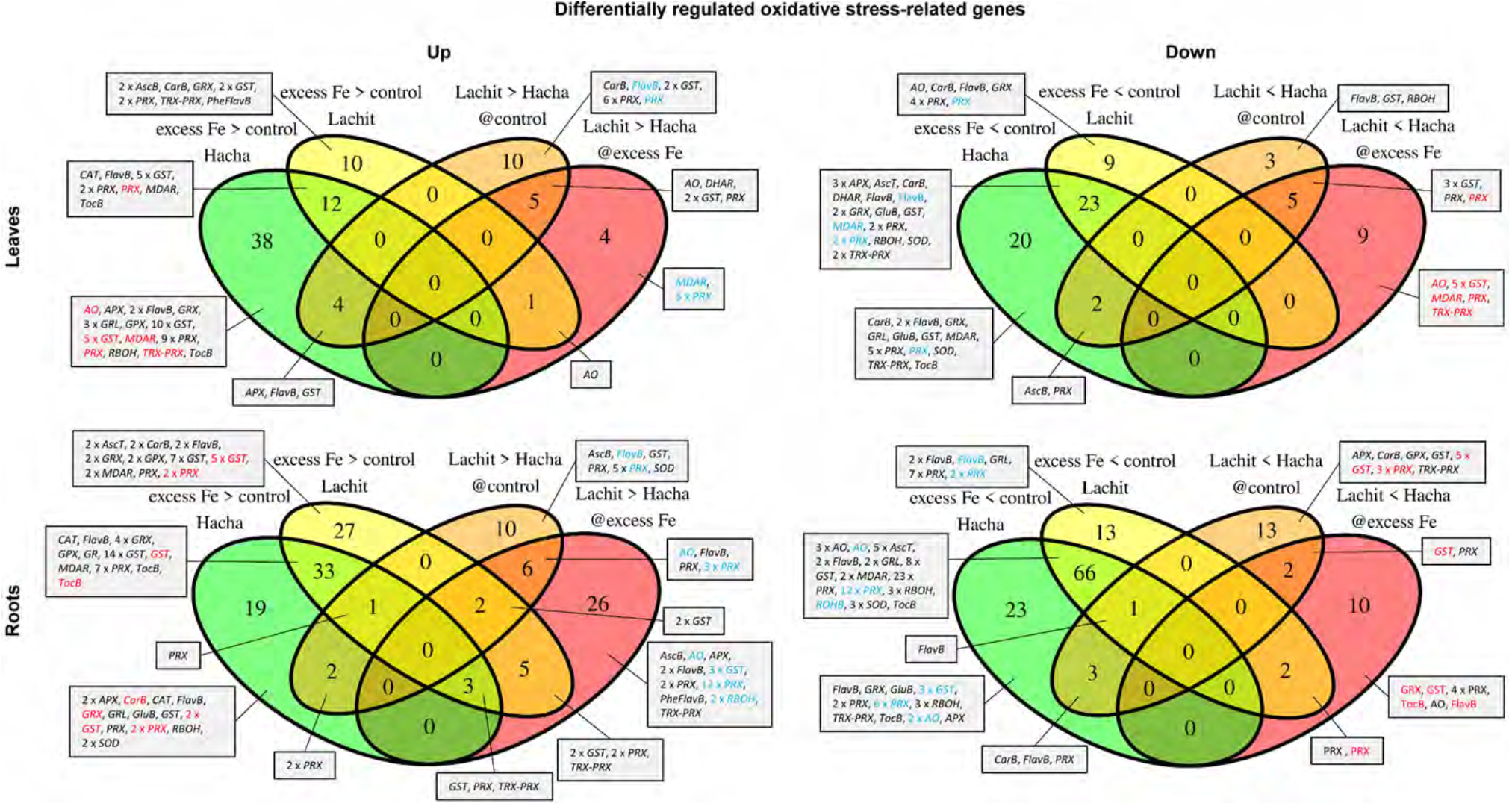
Four-way Venn diagrams of oxidative stress-related genes. Left, up-regulated genes (Up); right, down-regulated genes (Down); upper part, leaves; lower part, roots. The respective comparisons are indicated. Genes in blue, expressed at higher level in Lachit than in Hacha (left diagram) due to down-regulation in Hacha (right diagram). Genes in red, expressed at lower level in Lachit than in Hacha (right diagram) due to induction in Hacha (left diagram). Consequently, red and blue genes in the left diagram correspond to red and blue genes in the respective right diagram. Denoted are respective gene families or functions. Functions: AscB = ascorbate biosynthesis, AscT = ascorbate transport, CarB = carotenoid biosynthesis, FlavB = flavonoid biosynthesis, GluB = glutathione biosynthesis, PheFlavB = phenolics and flavonoid biosynthesis, TocB = tocopherol biosynthesis. Gene families: AO = ascorbate oxidase, APX = ascorbate peroxidase, CAT = catalase, DHAR = dehydroascorbate reductase, GPX = glutathione peroxidase, GRL = glutaredoxin-like, GRX = glutaredoxin, GST = glutathione-S-transferase, MDAR = monodehydroascorbate reductase, PRX = peroxidase, RBOH = respiratory burst oxidase, SOD = superoxide dismutase, TRX-PRX = thioredoxin-peroxidase. For more detailed information on the genes in this figure see Supplemental Tables S22 to S25.

